# PTE, a novel module to target Polycomb Repressive Complex 1 to the *Cyclin D2* oncogene

**DOI:** 10.1101/177097

**Authors:** Sarina R. Cameron, Soumyadeep Nandi, Tatyana G. Kahn, Juan I. Barrasa, Per Stenberg, Yuri B. Schwartz

## Abstract

Polycomb Group proteins are essential epigenetic repressors. They form multiple protein complexes of which two kinds, PRC1 and PRC2, are indispensable for repression. Although much is known about their biochemical properties, how PRC1 and PRC2 are targeted to specific genes is poorly understood. Here we establish the *Cyclin D2* (*CCND2*) oncogene as a simple model to address this question. We provide the evidence that coordinated recruitment of PRC1 and PRC2 complexes to *CCND2* involves a combination of a specialized PRC1 targeting element (PTE) and an adjacent CpG-island, which together act as a human Polycomb Response Element. Chromatin immunoprecipitation analysis of *CCND2* in different transcriptional states indicates that histone modifications produced by PRC1 and PRC2 are not sufficient to recruit either of the complexes. However, catalytic activity of PRC2 helps to anchor PRC1 at the PTE. Our analyses suggest that coordinated targeting of PRC1 and PRC2 complexes by juxtaposed AT-rich PTEs and CpG-islands may be a general feature of Polycomb repression in mammals.

## Introduction

Polycomb Group (PcG) proteins comprise a family of epigenetic repressors that prevent unscheduled transcription of hundreds of developmental genes (1–4). PcG proteins act in concert as multisubunit complexes. These are usually grouped into two evolutionary conserved classes called Polycomb Repressive Complex 1 (PRC1) and PRC2. Several auxiliary complexes, whose repertoire varies between species, further aid PRC1 and PRC2 to achieve robust repression (5).

PRC2 complexes contain a core with five subunits: EZH2 (or related protein EZH1), EED, SUZ12, RBBP4 (or closely related RBBP7) and AEBP2 (6,7), which together act as a histone methyltransferase specific to Lysine 27 of histone H3 (8–11). Tri-methylation of H3K27 (H3K27me3) at target genes is necessary for PcG repression (12). In addition to extensive tri-methylation of H3K27 at target genes, PRC2 complexes are also responsible for di-methylation of H3K27 within the entire transcriptionally inactive genome (13,14). This di-methylation happens via untargeted “hit-and-run” action and suppresses pervasive transcription (14).

The systematics of PRC1 group is more complicated as RING2 (or closely related RING1 protein), the central subunit of PRC1, are incorporated in large variety of protein complexes (Gao et al., 2012; Schwartz and Pirrotta, 2013). Here, for simplicity, we will use the PRC1 name for complexes that consist of one of the five variant chromodomain proteins (CBX2, CBX4, CBX6, CBX7 or CBX8), one of the three Polyhomeotic-like proteins (PHC1, PHC2 and PHC3), SCMH1 (or related SCML2) protein and a heterodimer between RING2 (or RING1) and one of the two closely related PCGF proteins: MEL18 (also known as PCGF2) and BMI1 (also known as PCGF4). These complexes, sometimes called canonical PRC1, can recognize H3K27me3 produced by PRC2 via the chromodomain of their CBX subunits (15–17). Mutations in genes encoding PRC1 subunits lead to embryonic lethality and miss-expression of *HOX* genes, indicating that PRC1 complexes are essential for PcG repression (18–21).

Other RING2/RING1 complexes contain RYBP (or closely related YAF2 protein) and one of the four PCGF proteins (PCGF1, PCGF3, PCGF5 and PCGF6). Complexes with different PCGF proteins have distinct additional subunits (5,22). None of these RING-PCGF complexes include CBX and PHC proteins and, therefore, cannot recognize the H3K27me3 mark. While PRC1 complexes are integral for PcG repression, the role of other RING-PCGF complexes, sometimes called non-canonical PRC1, is less clear. One of the complexes, which incorporates the RING2-PCGF1 dimer and the characteristic KDM2B subunit, is involved in PcG repression in mouse embryonic stem cells (23–25). However, some of the other RING-PCGF complexes have functions unrelated to PcG mechanisms (26).

PRC1 and other RING-PCGF complexes can act as E3 ligases *in vitro* to transfer a single ubiquitin group to Lysine 119 of histone H2A (H2AK119) (22,27,28). How the *in vivo* ubiquitylase activity of PRC1 differs from that of RYBP-containing RING-PCGF complexes is a subject of debate and there is disagreement in whether PRC1 is less catalytically active than its RYBP-containing counterparts (22,23,27). PRC2 is able to recognize H2AK119ub via its core subunit AEBP2 (29). However, the importance of H2AK119ub for PcG repression is not entirely clear. Thus mice, homozygous for the *Ring2* allele that encodes a catalytically inactive protein, show no missexpression of *Hox* genes and die much later than mice lacking PRC1 and PRC2 complexes (30). Likewise, *Drosophila*, which have H2AK118 ubiquitylation (analog of mammalian H2AK119ub) abolished by replacement of the corresponding H2A lysine to arginine, express *Hox* genes in proper manner and survive until late larval stages (31).

In *Drosophila*, where PcG system was first discovered and is the most studied, the targeting of PRC1 and PRC2 depends on designated Polycomb Response Elements (PREs). These approximately 1kb-long DNA elements are necessary and sufficient to recruit both complexes and tend to repress reporter genes when integrated elsewhere in the genome (32). The ability of PREs to repress is not absolute. If a target gene gets activated at early embryonic stages, the de-repressed state is often inherited by the progenitor cells (33) and the binding of PcG complexes and H3K27 trimethylation is impaired (34–37).

PRC1 can specifically recognize the H3K27me3 modification catalyzed by PRC2 and PRC2 can recognize H2AK119ub produced by PRC1. However, neither histone modification is necessary or sufficient to target PRC complexes to PREs (14,38). Instead, multiple lines of evidence indicate that PREs contain combinations of recognition sequences for different DNA binding factors. These factors act cooperatively to recruit PRC1 and PRC2, which themselves cannot bind DNA in sequence specific fashion (reviewed in: (32)). Our recent work indicates that PRC1 can bind PREs in the absence of PRC2. However, at many PREs, PRC2 requires PRC1 for binding (38). Consistently, in cells where PcG target genes are active they often lose H3K27me3 and PRC2 but retain significant amount of PRC1 bound at PRE sites (34,36,37).

PRC1 and PRC2 complexes are evolutionary conserved and target many of the same genes in *Drosophila* and mammals (4). It is, therefore, hard to imagine that mechanisms to bring PRC complexes to these genes have evolved completely independently. The question, therefore, is which parts of the targeting process are similar and which have diverged since flies and mammals split from their common ancestor several hundred million years ago. A number of mouse and human DNA fragments were reported to resemble PREs in their ability to recruit either PRC1 or PRC2 complexes when integrated elsewhere in the genome (39–46). Yet, the overall picture remains obscure. First, in contrast to paradigmatic *Drosophila* PREs, few of the reported mammalian PRE-like elements were mapped to high precision or their recruitment properties confirmed by independent investigations. Second, some of these elements were said to recruit both PRC1 and PRC2 while others appear to recruit only one of the complexes. Lastly, little is known about the mechanisms implicated in targeting.

Several independent investigations concur that DNA sequences with high density of unmethylated CpG di-nucleotides (so-called CpG islands) are sufficient to recruit PRC2 but not PRC1 in mouse embryonic stem cells (41–43). Presence of binding sites for transcription factors REST and SNAIL within a CpG island may further help PRC2 recruitment (39,47). Studies from Klose and Brockdorff groups put forward a model linking CpG-islands to PRC2. According to this model, mouse Ring2-Pcgf1-Kdm2B complex recognizes unmethylated CpG-islands via the CXXC domain of the Kdm2B subunit. Once recruited to a CpG island the complex mono-ubiquitylates surrounding nucleosomes at H2AK119 which, in turn, serve as a binding platform for PRC2 (23,48). Clearly different from how PRC2 is recruited in *Drosophila* (14,38), the model is supported by observations that tethering a minimal catalytically active H2AK119 ubiquitylase is sufficient for PRC2 recruitment (23). Yet, the central role of H2AK119ub implied by the model, is at odds with reports that H2AK119ub is dispensable for early mouse development (30). Furthermore, which DNA elements, if any, target PRC1 complexes and whether H3K27 methylation is involved in the process remains an open question.

Here we establish the *Cyclin D2* (*CCND2*) oncogene as a simple model system to address these questions. Using this system, we find that coordinated recruitment of PRC1 and PRC2 complexes to *CCND2* involves a combination of a specialized PRC1 targeting element (PTE) and an adjacent CpG-island, which we view as a functional analogue of *Drosophila* PREs. Chromatin immunoprecipitation analysis of *CCND2* in different transcriptional states suggest that H2AK119ub and H3K27me3 do not serve as binding platforms for either complex. Yet, catalytic activity of PRC2 helps PRC1 to bind the PTE. Further analyses suggest that coordinated targeting of PRC1 and PRC2 complexes by juxtaposed PTEs and CpG-islands is not unique to the *CCND2* gene and that PRC1 targeting by compact DNA elements may be an ancestral feature of Polycomb repression.

## Results

PRC1 and PRC2 complexes usually act together to effect epigenetic repression. However, in experiments with cultured *Drosophila* cells, we noted PcG target genes that retained PRC1 at PREs in the absence of PRC2 and H3K27me3 when they happened to be transcriptionally active (36,37). This fueled our interest to a region approximately 4.8kb upstream of the Transcription Start Site (TSS) of the human *Cyclin D2* (*CCND2*) gene. Re-analyzing previously published Chromatin Immunoprecipitation (ChIP) binding profiles (49), we noticed that in the embryonic teratocarcinoma NT2-D1 cells this region is strongly immunoprecipitated with antibodies against BMI1 and MEL18 but very weakly with antibodies against EZH2 and H3K27me3 (Figure 1A-C). This is in stark contrast to all other sites on human chromosomes 8, 11 and 12, interrogated in this experiment, which show strong precipitation with anti-EZH2 and anti-H3K27me3 antibodies whenever they are strongly precipitated with antibodies against BMI1 or MEL18 (Figure 1A, C).

**Figure 1.**
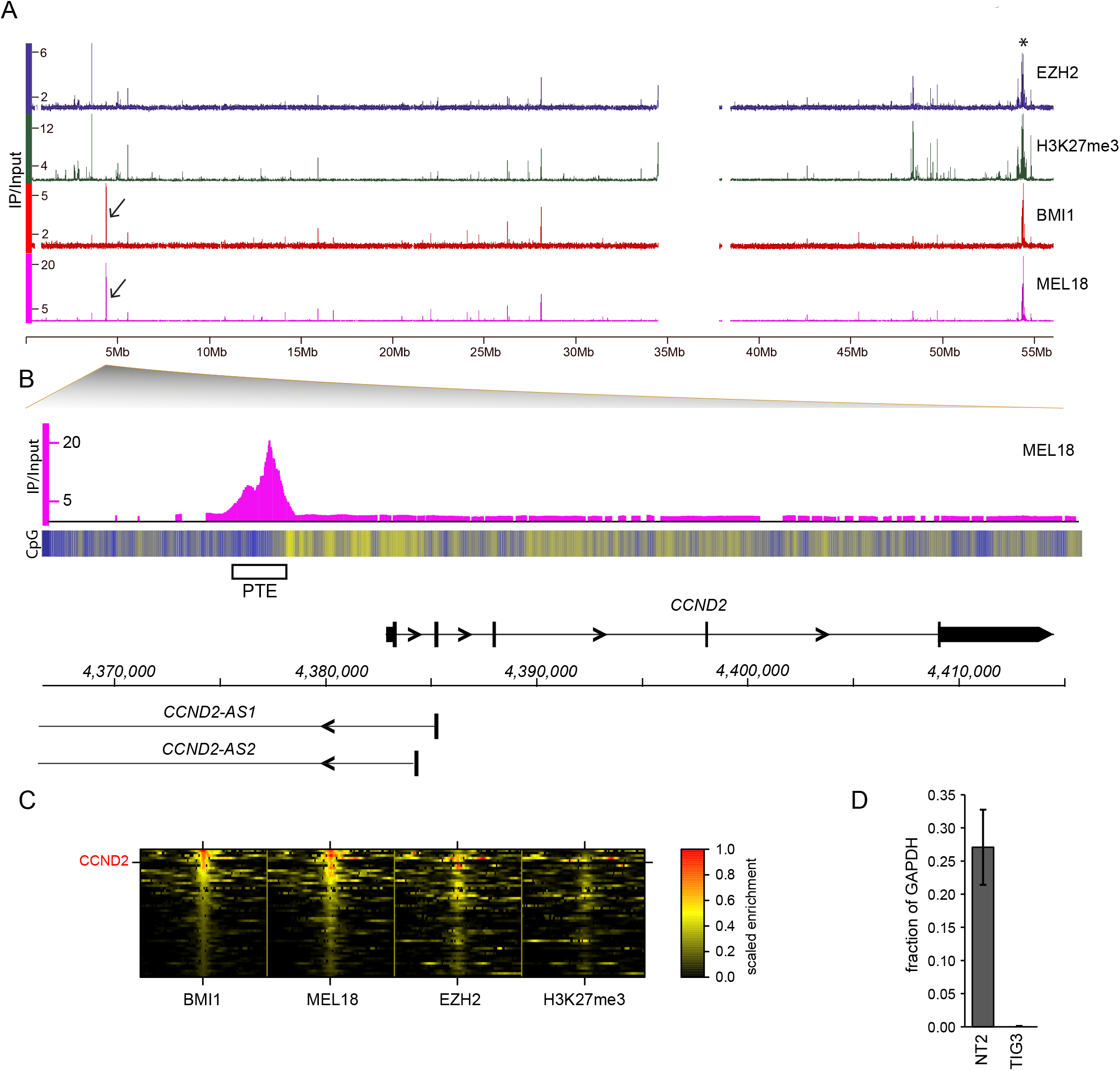
Disparate binding of PRC1 and PRC2/H3K27me3 reveals putative PRC1 targeting element upstream of the *CCND2* gene. **A**. ChIP-on-chip profiles of MEL18, BMI1, H3K27me3 and EZH2 along the 55Mb of the left half of the human Chromosome 12. The gap corresponds to repetitive sequences around the centromere. Asterisk indicates the position of the *HOXC* cluster, the classic PcG target. Arrows mark strong anti-MEL18 and anti-BMI1 ChIP signals at the *CCND2* locus. **B**. Close up view of MEL18 ChIP profile at the *CCND2* locus. The heat-map underneath MEL18 binding profile shows the fraction of CpG nucleotides within 100bp sliding window (ranging from dark blue - lowest to bright yellow - highest). The DNA sequence underneath MEL18 peak (white rectangle, PTE) is CpG poor but immediately adjacent is the CpG-rich region that extends towards the *CCND2* TSS. The *CCND2* gene is transcribed from left to right. The lncRNAs *CCND2-AS1* and *CCND2-AS2* are transcribed from right to left. The scale indicates genomic positions in Hg19 coordinates. **C**. With an exception of *CCND2*, genomic regions that show strong ChIP-chip signals for MEL18 and BMI1 are also strongly precipitated with anti-EZH2 and H3K27me3 antibodies. The heat-map plots show ChIP-chip signals over 10kb regions centered on BMI1 ChIP signal peaks. Each row represents one region and these are sorted by the BMI1 ChIP signal strength. The enrichment values for each protein are scaled such that the strongest genomic binding corresponds to 1. **D**. RT-qPCR measurements indicate that *CCND2* is highly expressed in NT2-D1 (NT2) but very little in TIG-3 cells. Histograms show the average and the scatter (whiskers) between two independent experiments. Values are normalized to the expression of the housekeeping *GAPDH* gene.

Quantitative RT-PCR measurements show that, like in the *Drosophila* cases, the *CCND2* gene of NT2-D1 cells is highly transcribed (Figure 1D). In contrast, in TIG-3 embryonic fibroblasts the *CCND2* gene is transcriptionally inactive (Figure 1D). In these cells, it is decorated with H3K27me3, binds CBX8 and SUZ12 and gets upregulated upon the knockdown of EZH2, EED, SUZ12 and BMI1 (50). This indicates that *CCND2* is an ordinary PcG target gene which, when transcriptionally inactive, acquires the chromatin state characteristic of PcG repression but binds a lot of PRC1 and little PRC2 and H3K27me3 in cells where it is transcriptionally active. This finding is significant in three ways. First, it shows that PRC1 can bind target genes when very little H3K27me3 is present. Second, it suggests that, like in *Drosophila*, the binding of PRC1 in the vicinity of a gene is not by itself repressive. Third, it points to a location of a DNA element that may be responsible for PRC1 recruitment to *CCND2* (putative PRC1 Targeting Element). It also suggests that *CCND2* is as a good model to investigate how the targeting of human Polycomb complexes is coordinated.

### PRC1 and PRC2 binding to the *CCND2* gene in alternative chromatin states

With this in mind, we performed quantitative ChIP analysis of PRC1, PRC2 and H3K27me3 binding at the *CCND2* gene in the NT2-D1 and TIG-3 cell lines. To select informative protein targets for immunoprecilitation, we first measured the mRNA levels for genes encoding PRC1 subunits to know which of the alternative variants are available for interrogation in each of the cell lines (Figure S1). As shown by RT-qPCR, *MEL18* mRNA is highly abundant in NT2-D1 cells and slightly less abundant in TIG-3 cells (Figure S1). The *BMI1* mRNA is present at moderate level in both NT2-D1 and TIG-3 cells. *RING1* mRNA level is low in both cell lines but *RING2* mRNA is highly abundant in NT2-D1 cells and at medium level in TIG-3 cells (Figure S1). Out of five *CBX* genes implicated in PcG regulation (5), mRNA levels for *CBX4*, *CBX6* and *CBX7* are low in both NT2-D1 and TIG-3 cells. *CBX8* mRNA is at the edge of the detection in both cell lines and *CBX2* mRNA is highly abundant in NT2-D1 but barely detectable in TIG-3 cells.

Immunoprecipitations of the formaldehyde crosslinked NT2-D1 chromatin with antibodies against MEL18, BMI1, CBX2 and RING2 give essentially the same results (Figures 2, S2). The ChIP signals peak within a putative PRC1 Targeting Element (PTE) and recede steeply at both sides to reach a background level half way between the PTE and the *CCND2* TSS. In TIG-3 cells, where *CCND2* is transcriptionally inactive, the putative PTE remains the strongest precipitated site with ChIP signals similar to those detected in NT2-D1 cells. However, in these cells, in addition to the PTE the entire upstream region of the *CCND2* gene, including the TSS, is also immunoprecipitated, albeit at a much lower level. In contrast to PRC1, ChIPs with antibodies against SUZ12 and H3K27me3 give enrichment profiles that differ dramatically between the two cell lines. In NT2-D1 cells, SUZ12 ChIP signals are very weak, compared to that at the positive control *ALX4* gene, barely raising above the background at the PTE (Figure 2). These are paralleled by weak ChIP signals for H3K27me3. In contrast, in TIG-3 cells, ChIP signals for both antibodies are very strong and on par with those at *ALX4* gene. Strikingly, their profiles do not match those for PRC1. Instead, the SUZ12 and H3K27me3 profiles are broad and shifted away from the CpG-poor PTE into the adjacent CpG-rich region (Figures 2, 1B). The obvious difference between the PRC1 and SUZ12/H3K27me3 ChIP profiles is instructive. First, it indicates that a high level of H3K27me3 is not sufficient to recruit PRC1 to an extent seen at the putative PTE and that other mechanisms must contribute to PRC1 recruitment to this element. Second, it suggests that the putative PTE is not capable to recruit PRC2 as efficiently as activities linked to the adjacent CpG-rich region.

**Figure 2.**
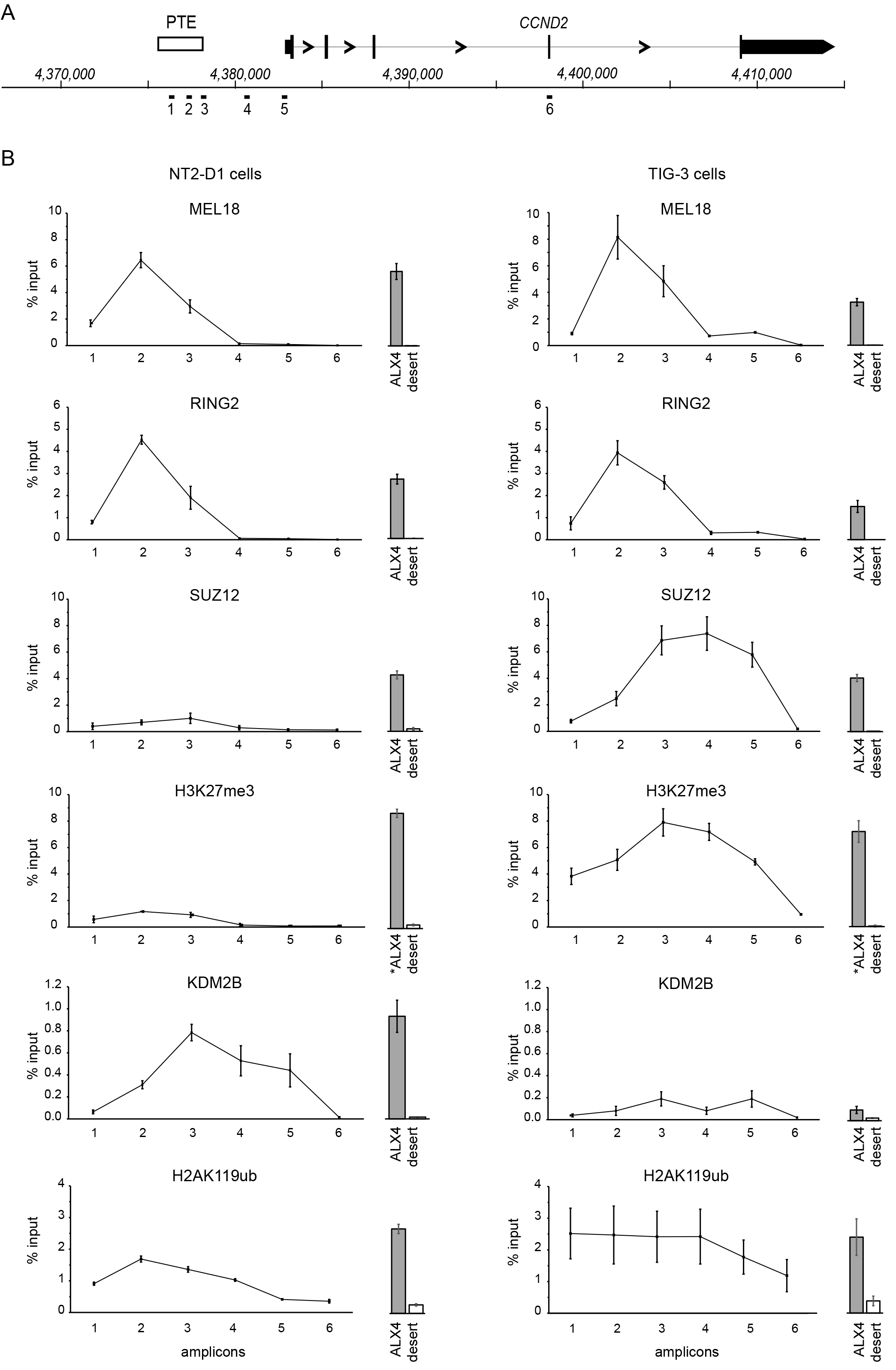
Polycomb proteins and associated histone modifications at the active and repressed *CCND2* gene. **A**. The schematic of the *CCND2* locus. Positions of PCR amplicons analyzed in **B** are shown below the coordinate scale. Note that amplicon N2 corresponds to the summit of MEL18 peak and amplicon N5 corresponds to the *CCND2* TSS (see: Tables S4, S5 for sequence information). **B**. ChIP profiles of MEL18, RING2, SUZ12, H3K27me3 KDM2B and H2AK119ub in NT2-D1 (left column) and TIG-3 (right column) cells. The immunoprecipitation of ALX4 gene repressed by PcG in NT2-D1 and TIG-3 cells was used as positive control. The immunoprecipitation of a gene desert region (Chr.12: 127711096127713493; Hg19) was used as a negative control. Histograms for both are shown to the right of the each graph and are to the same y-axes scale. ALX4* positive control region for H3K27me3 ChlPs is offset from the ALX4 region used for the profiling of other proteins. This is done to account for relatively low H3K27me3 at the site of the highest PcG occupancy overlapped by ALX4 amplicon. All histograms and graphs show the average and the scatter (whiskers) between two independent experiments.

### KDM2B, H2AK119 ubiquitylation and recruitment of PRC2

Mouse Ring2-Pcgf1-Kdm2B complex can bind unmethylated CpG islands and catalyze the monoubiquitylation of H2AK119 (24,25,51), which was proposed to directly recruit PRC2 (23). According to data from the ENCODE project, in NT2-D1 cells the CpG-region upstream of the *CCND2* TSS is unmethylated (52). We therefore asked to what extent the apparent strong binding of PRC2 over the CpG-rich region in TIG-3 cells is linked to the presence of KDM2B-RING2 complexes and H2AK119ub.

KDM2B is not crosslinked to chromatin with formaldehyde (24), so we used a combination of formaldehyde and EGS (ethylene glycol bis(succinimidyl succinate)) to prepare chromatin for KDM2B mapping. Somewhat surprisingly, the KDM2B ChIP signals are higher in NT2-D1 cells, probably because in these cells the *KDM2B* gene is expressed much stronger (Figure S1), but are qualitatively similar to those in TIG-3 cells (Figure 2). In contrast to PRC1, the KDM2B ChIP profile is shifted from the putative PTE into adjacent CpG-rich DNA. In NT2-D1 cells, most of this region (amplicons 4 and 5) is not precipitated with the anti-RING2 antibodies regardless of whether we perform RING2 ChIP from chromatin crosslinked with formaldehyde and EGS or formaldehyde alone (Figure S3). It appears that when *CCND2* is transcriptionally active KDM2B binds the CpG-island independently of RING2. Consistently, in NT2-D1 cells the H2AK119ub ChIP signal peaks at the putative PTE and recedes to the background level between the PTE and the *CCND2* TSS (Figure 2). In TIG-3 cells, the region precipitated with anti-H2AK119ub antibodies is broad and coextensive with that precipitated with anti-SUZ12 and anti-H3K27me3 antibodies. The strength of the H2AK119ub ChIP signal in TIG-3 cells is comparable to that over the putative PTE in NT2-D1 cells. This suggests that either H2A119ub is not by itself sufficient to recruit much PRC2 and additional processes must be in place for efficient recruitment, or that something prevents H2A119ub-mediated PRC2 recruitment when *CCND2* is transcriptionally active.

### *CCND2* PTE can autonomously recruit PRC1 and is evolutionary conserved

To investigate whether the DNA fragment underneath the *CCND2* PRC1 binding peak is indeed a PRC1 targeting element, we tested whether it can autonomously recruit PcG complexes when integrated elsewhere in the genome. To this effect, we cloned the 2.4kb fragment covering the putative PTE into a lentiviral vector (Figure 3A) and integrated it back into the genome of NT2-D1 cells by viral transduction. To distinguish transgenic and endogenous copies of the putative PTE we identified a small stretch of nucleotides close to the summit of the MEL18/BMI1 peak that shows little conservation within mammalian species and substituted five of those nucleotides in the transgenic copy to create an annealing site for a specific PCR primer (Figure S4). An analogous construct containing a 2.4kb fragment from a gene desert region on chromosome 12 that showed no binding of PcG proteins and H3K27me3 (Figure S4) and an empty vector were integrated in parallel as negative controls. The transduced cell lines were genotyped by PCR to validate their identity (Figure 3A, Figure S5).

**Figure 3.**
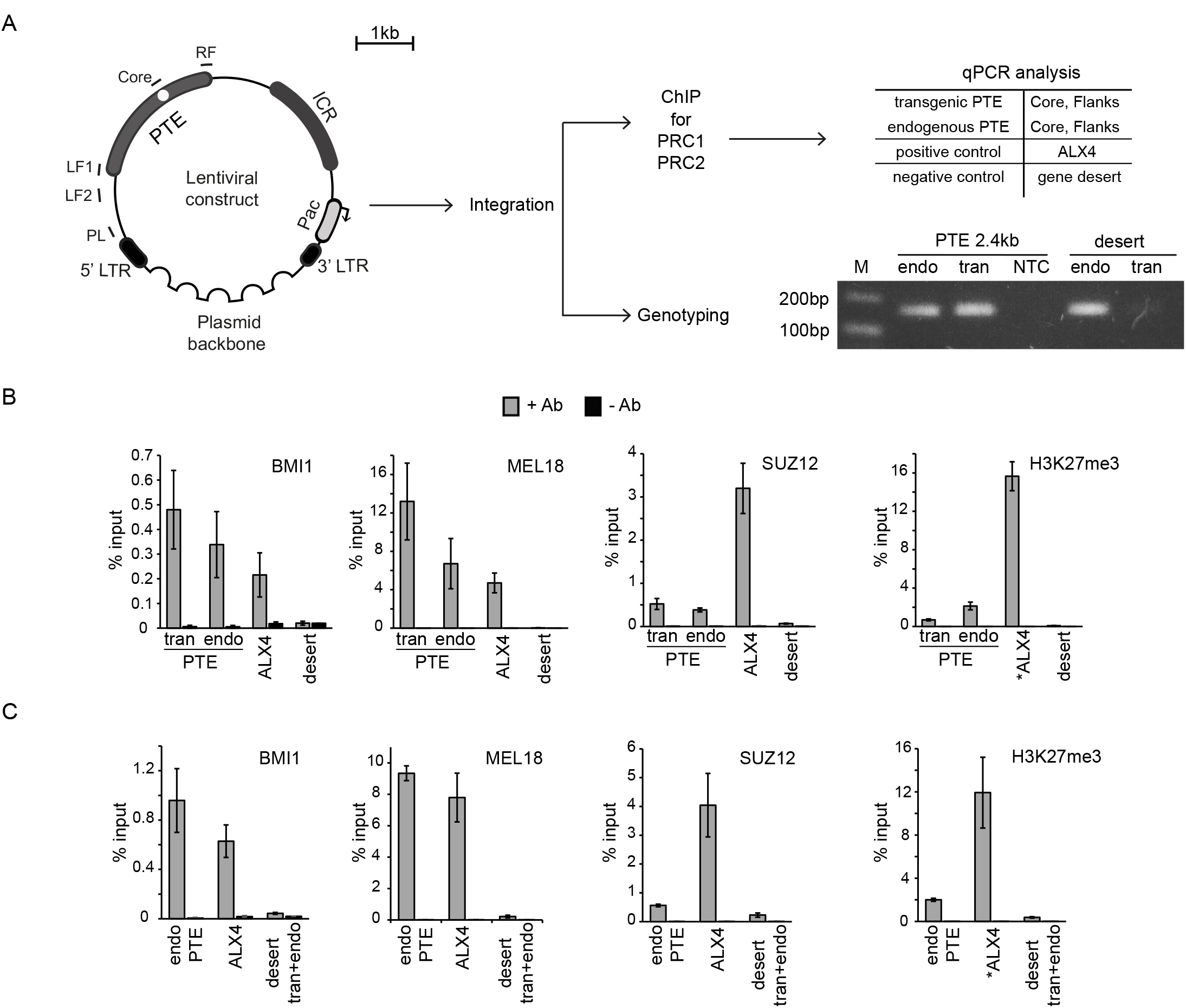
The *CCND2* PTE can autonomously recruit PRC1. **A**. The schematic of experimental setup. Tested fragments, represented by 2.4kb PTE fragment, were cloned in the lentiviral vector. The vector contained *Pac* puromycin resistance gene shielded from potential repression by the chromatin insulator from mouse *Igf2/H19* Imprinting Control Region. The position of an engineered annealing site for transgene-specific PCR primer is indicated by white circle. Positions of additional amplicons are indicated as short lines. Following viral transduction and puromycin selection, cells were genotyped and collected for ChIP-qPCR assays. The results of a representative analysis of PCR products shows that transgene-specific primers amplify the product of expected size from genomic DNA of cells transduced with putative 2.4kb PTE but not with a control (gene desert) construct (see Figure S4 for additional details). Comparison of immunoprecipitations of the 2.4kb PTE fragment (**B**) and the control 2.4kb fragment from Chromosome 12 gene desert (**C**) indicates that the PTE 2.4kb robustly recruits PRC1 and little PRC2 or H3K27me3. The precipitation of transgenic (tran) and endogenous (endo) PTE fragments was assayed using corresponding “Core” amplicons. The immunoprecipitations of the ALX4 and Chromosome 12 gene desert regions were assayed in parallel as positive and negative controls. The PCR amplicon used to assay the immunoprecipitation of the gene desert fragment in (**C**) does not discriminate between transgenic and endogenous copies. Since the endogenous site does not bind PcG any elevated ChIP signal would come from the transgenic copy. All histograms show the average and the scatter (whiskers) between two independent experiments.

As summarized in Figure 3B, the 2.4kb transgenic PTE fragment is immunoprecipitated with antibodies against BMI1 and MEL18. The precipitation is robust and as strong as that of the endogenous PTE upstream of *CCND2*. This is in contrast to the transgenic insertion of the negative control fragment from the gene desert (Figure 3C) or the empty vector (Figure S6) whose precipitation is very low and close to the ChIP background. The immunoprecipitation of the transgenic 2.4kb fragment with the antibodies against SUZ12 and H3K27me3 is weak and comparable to that of the endogenous site (Figure 3B). Taken together these observations suggest that the 2.4kb DNA fragment underneath the *CCND2* PRC1 binding peak indeed contains an element capable of autonomous recruitment of PRC1 complexes.

The *CCND2* PTE is evolutionary conserved suggesting that its function is important. Thus, mouse *Ccnd2* resides within large 32Mbp block of shared synteny between mouse chromosome 6 and human chromosome 12. Aside of the open reading frame, which has high DNA sequence conservation, the 10kb sequence upstream of the *Ccnd2* TSS contains multiple blocks predicted as evolutionary conserved DNA elements (Figure 4A). One of these blocks corresponds to ~300bp sequence that is directly underneath the PRC1 binding peak within the human *CCND2* PTE.

**Figure 4.**
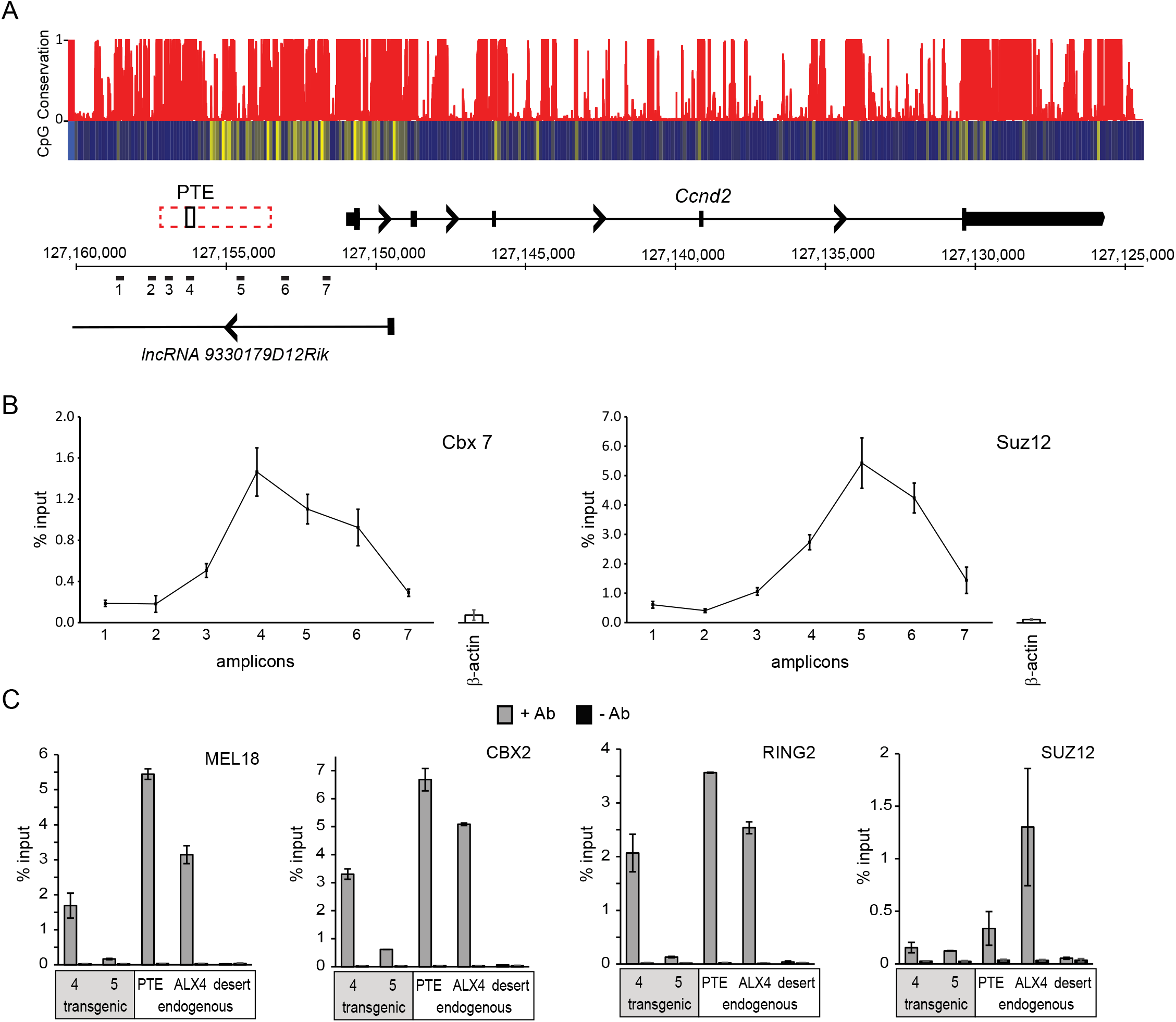
Mouse *Ccnd2* locus contains a PTE. **A**. The schematic of the mouse *Ccnd2* locus. PhastCons sequence conservation scores based on the alignment of 60 vertebrate genomes illustrate that putative mouse PTE is an evolutionary conserved element. The heat-map underneath the conservation score graph shows the number of CpG nucleotides within 100bp sliding window (ranging from dark blue=0 to bright yellow=10). Similar to the human *CCND2* locus, the CpG-poor PTE is adjacent to a CpG-island that extends towards the *Ccnd2* TSS. The *Ccnd2* gene is transcribed from left to right. Also shown is lncRNA (*9330179D12Rik*) of unknown function. Positions of PCR amplicons analyzed in **B** and **C** are shown below the scale in mm10 genomic coordinates. Dashed rectangle indicates the 3.6kb fragment tested for autonomous recruitment of PRC1 in human cells. Black rectangle marks position of evolutionary conserved DNA sequence that corresponds to ~300bp fragment directly underneath the PRC1 binding peak within the human *CCND2* PTE. **B**. Cbx7 and Suz12 ChIP profiles across upstream region of *Ccnd2* in the mouse F9 cells. Note that Suz12 profile is offset from the PTE and the Cbx7 peak into the CpG-rich region. The immunoprecipitation of *β-actin* gene (shown as bars to the right of the each graph) was used as negative control. Here and in **C** bar plots and graphs are to the same scale and show the average and the scatter (whiskers) between two independent experiments. **C**. Chip analysis of 3.6kb mouse PTE fragment integrated in the genome of human NT2-D1 cells shows that it robustly recruits PRC1 but little PRC2. Amplicons 4 and 5 shown in **A** were used to assay the transgenic mouse fragment. Immunoprecipitation of the endogenous human *CCND2* PTE and the *ALX4* locus were assayed in parallel as positive controls and precipitation of the Chromosome 12 gene desert was assayed as negative control.

In mouse F9 testicular teratoma cells, where the *Ccnd2* gene is transcriptionally inactive, its upstream region is immunoprecipitated with antibodies against Cbx7 and Suz12 (Figure 4B). This suggests that the mouse *Ccnd2* is a PcG target gene. Similar to what is seen at inactive human *CCND2* in TIG-3 cells, the ChIP signal for Cbx7 is highest at the site corresponding to the evolutionary conserved block within the PTE but the highest Suz12 ChIP signal is shifted from the Cbx7 peak towards the TSS into the CpG-rich area. To extend this comparison further, we cloned the 3.6kb fragment covering the putative mouse PTE into the same lentiviral vector used to test the human PTE (Figure 4A) and integrated it into the genome of the human NT2-D1 cells by lentiviral transduction. ChIP-qPCR analysis indicates that the evolutionary conserved central part of the mouse 3.6kb fragment is robustly precipitated with antibodies against MEL18, CBX2 and RING2 and weakly with antibodies against SUZ12 (Figure 4C). This indicates that the mouse *Ccnd2* gene is also equipped with a PTE.

### The *CCND2* PTE is modular and its activity depends on specific DNA sequences

*Drosophila* PREs, the closest analogues of the *CCND2* PTE, contain recognition sequences for multiple unrelated DNA binding proteins that cooperate to provide robust PcG repression. A typical PRE has a size of about 1kb. Often such PREs may be subdivided into fragments of a few hundred base pairs that can still recruit PcG proteins in transgenic assays but the recruitment is less robust (53–55). We, therefore, wondered whether the *CCND2* PTE is smaller than the 2.4kb fragment tested in our initial transgenic assay and whether the PTE contains a single core recruiting element or multiple weaker elements that cooperate. To address these questions we examined PRC1 recruitment by sub-fragments of the 2.4kb *CCND2* PTE (Figure 5A). ChIP analysis of PRC1 binding in cells transduced with corresponding lentiviral constructs indicates that the 1kb fragment (PTE 1.2) centered on the summit of the MEL18 and BMI1 binding peak and the larger overlapping fragments (PTE 1.1 and PTE 1.3) recruit MEL18, BMI1and CBX2 as efficiently as the full-length 2.4kb fragment or the endogenous *CCND2* PTE (Figure 5B-D). In contrast, precipitation of the PTE 1.2 fragment with anti-SUZ12 antibodies is weak (Figure 5B) reinforcing our conclusion that the *CCND2* PTE is not an efficient PRC2 recruiter.

**Figure 5.**
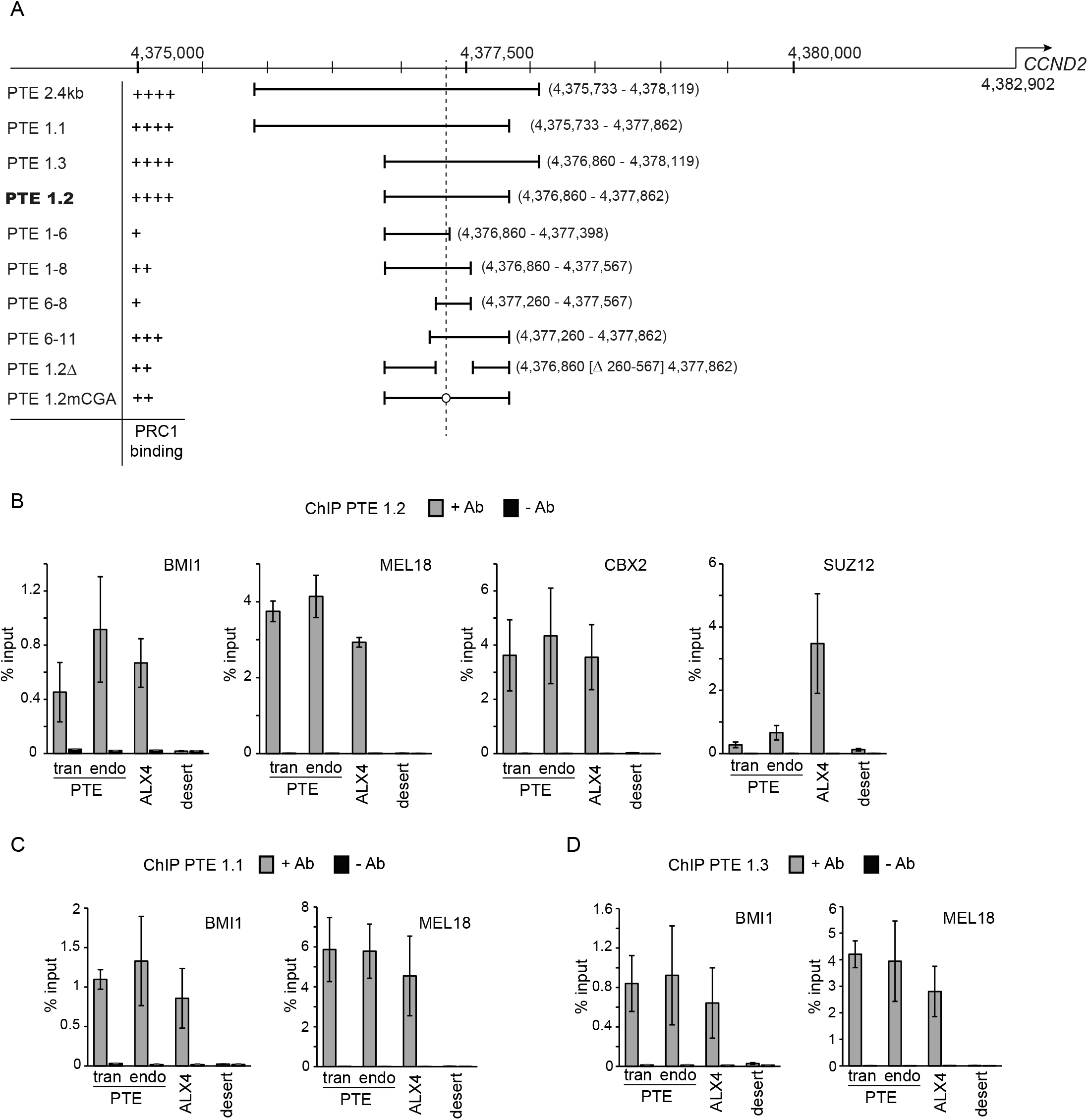
Transgenic dissection of the *CCND2* PTE. **A**. Schematic representation of PRC1 recruitment capacity of different fragments from the *CCND2* PTE region. “++++” indicates recruitment comparable to endogenous locus, “+++” indicates robust recruitment but weaker than at endogenous locus, “++” indicates intermediate recruitment, “+” indicates weak recruitment. The fragment positions are shown in relation to *CCND2* TSS and Hg19 genomic coordinates. The minimal 1kb PTE 1.2 fragment capable of robust PRC1 recruitment is indicated in bold. The PTE 1.2mCGA construct corresponds to PTE 1.2 fragments with mutated “CGA” motif. **B**. ChIP-qPCR analysis of the PTE 1.2 transgene. The precipitation of transgenic DNA (tran) is compared to that of the endogenous *CCND2* PTE region (endo), unrelated PcG repressed gene ALX4 and the gene desert region (desert). Here and in **C**, **D** histograms show the average and the scatter (whiskers) between the two independent experiments. ChIP-qPCR analyses of transgenes containing PTE 1.1 (**C.**) and PTE 1.3 (**D.**) indicate that both fragments bind PRC1 to an extent seen at the endogenous *CCND2* PTE.

Further dissection indicates that smaller sub-fragments of PTE 1.2 cannot recruit PRC1 as efficiently as the full-length fragment (Figures 6, S7). When integrated elsewhere in the genome, the left (PTE 1-6) or the central (PTE 6-8) parts are immunoprecipitated with anti-MEL18 antibodies very weakly (Figure 6B-C) and their precipitation with anti-BMI1 antibodies is at the edge of detection (Figure S7). This is likely, because in NT2-D1 cells BMI1 is less abundant than MEL18 (Figure S1). The immunoprecipitation of larger fragments that combine the left and the central parts (PTE 1-8) or the central and the right parts (PTE 6-11) is more efficient (Figures 6D-E, S7). Consistent with the asymmetric shape of the MEL18/BMI1 ChIP-chip peak (Figure 1B), the PTE 6-11 fragment shows the strongest immunoprecipitation from all PTE 1.2 sub-fragments tested. This suggests that the central and the right parts of the 1kb PTE have greater contribution to PRC1 recruitment. Importantly, the synthetic fragment (PTE1.2Δ) that includes both the left and the right parts of PTE 1.2 but lacks the central part is still immunoprecipitated (Figure 6F, S7). Overall, these results indicate that the *CCND2* PTE consists of at least three separable modules, all of which contribute to the recruitment with the central and the right modules being more important.

**Figure 6.**
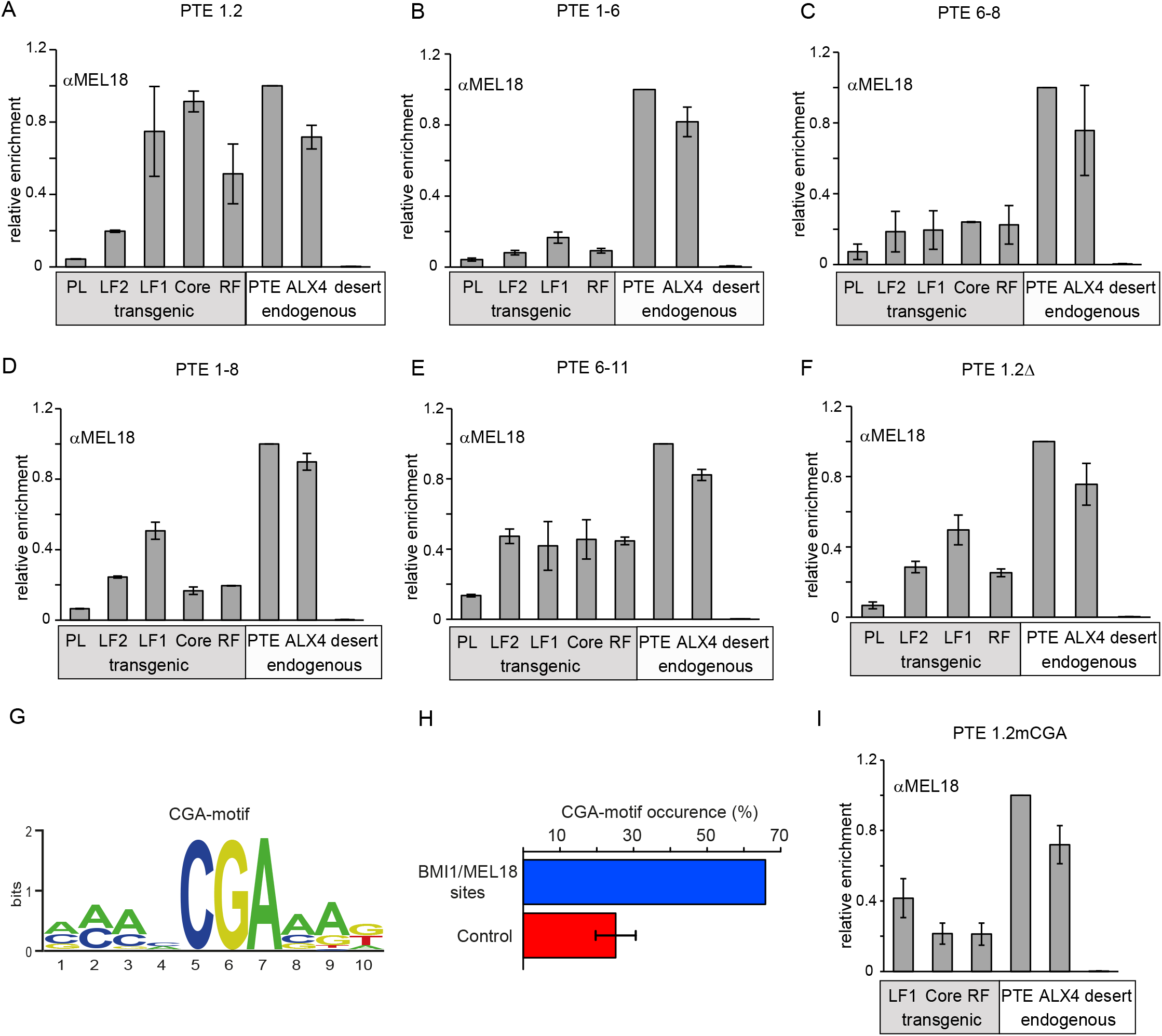
Multiple modules and the “CGA” motif contribute to PRC1 recruitment by the *CCND2* PTE. **A.-F.** ChIP analysis of MEL18 binding to a set of transgenic constructs spanning different parts of PTE 1.2 indicates that it consists of at least three separable modules, all of which contribute to the recruitment but the central and the right modules contributing more. Positions of various amplicons along the transgenic constructs (grey box) are illustrated in Figure 3A. Note that some transgenic constructs lack some of the amplicons. Precipitation of transgenic fragments is compared to internal positive and negative controls (white box). The histograms show average ChIP yields normalized to that of the endogenous *CCND2* PTE (amplicon 2) with whiskers indicating the scatter between two independent experiments. **G.** Logo representation of the “CGA”-motif. **H.** The occurrence of the “CGA”-motif within high confidence genomic BMI1/MEL18 bound sites (blue) is much higher than that within control regions (red). The whiskers mark 95% confidence interval of the mean based on hundred permutations of the control group. **I.** ChIP analysis of MEL 18 binding to transgenic 1kb PTE 1.2 fragment with the mutated “CGA”-motif Four-fold reduction of ChIP signals suggests that the “CGA”-motif is important for PRC1 recruitment by the PTE.

Both sequence specific DNA binding activity and *cis*-acting non-coding RNAs (ncRNAs) have been implicated in the recruitment of mammalian PcG proteins (56). Recent genome annotation indicates two long non-coding RNAs CCND2-AS1 and CCND2-AS2, which originate within the second exon or the first intron of *CCND2* (Figure 1B). They are transcribed in the opposite direction to *CCND2* and traverse over the PTE to terminate some 27kb away from their TSS (57). These lncRNAs are unlikely to play a critical role in autonomous recruitment by *CCND2* PTE as their transcription start sites were not included in our lentiviral constructs. We, therefore, sought evidence of a specific DNA binding activity targeting the PTE 1.2 fragment. The analysis of the PTE 1.2 DNA sequence revealed no potential binding sites for YY1, RUNX1/CBFβ, REST and SNAIL (Figure S8A), the sequence specific DNA binding proteins previously implicated in the recruitment of mammalian PcG proteins (39,46,47,58). Consistently, we and others detect no YY1 binding to the *CCND2* PTE (49,52). Mining the ENCODE data shows that the PTE 1.2 fragment does not bind any of the transcription factors mapped to date (Figure S8A) although multiple transcription factors bind the adjacent CpG-rich region.

Since we did not find any binding sites for known transcription factors, we analyzed the *CCND2* PTE for sequences that might bind proteins not yet implicated in PcG targeting. We hypothesized that some of the other PcG target genes, represented by high-confidence BMI1/MEL18 binding sites on the three human chromosomes surveyed by ChIP (49), may have elements similar to the *CCND2* PTE. In this case, they may use some of the same DNA binding proteins for PcG recruitment and may be distinguished by the same sequence motifs. To test this conjecture, we applied multivariate modelling (59,60) and searched for all possible 6 nucleotide long sequence words predictive of high-confidence MEL18/BMI1 binding sites compared to 1000 random sequences not bound by MEL18 or BMI1 and matched for the distance to the closest TSS. This approach yielded two motifs that we dubbed “CGA” and “CGCG” (Figure 6G, S8B). While the “CGCG”-motif is likely an outcome of the close proximity between PTEs and CpG-islands, the “CGA”-motif is interesting. First, it is present right at the summit of the *CCND2* MEL18/BMI1 peak and its position within the *CCND2* PTE is highly evolutionarily conserved (Figure S8C). Second, the “CGA”-motif is significantly enriched in high-confidence MEL18/BMI1 binding sites (Figure 6H). Overall, these observations suggest that the “CGA”-motif is a common feature of MEL18/BMI1 bound sites and may be involved in targeting PRC1 to *CCND2* PTE.

To test the role of the “CGA”-motif for recruitment by the *CCND2* PTE we generated a lentiviral construct that contained the 1kb PTE 1.2 fragment in which the CG dinucleotide within the “CGA”-motif was replaced with AA. ChIP-qPCR analysis of transduced NT2-D1 cells shows that antibodies against PRC1 components precipitate the mutated PTE1.2 fragment (PTE1.2mCGA) four-fold weaker than the wild-type variant (Figure 6I, S9). The reduction of ChIP signals is comparable to that seen after deleting the entire central part of the PTE (PTE1.2Δ construct). This suggests that the “CGA”-motif makes an important contribution to the recruitment by the evolutionary conserved central part of the PTE. The loss of ChIP signals upon mutation of the “CGA”-motif or deletion of the central part of the PTE is significant but not complete. This indicates that other sequences within the 1kb PTE contribute to recruitment. Consistently, Electrophoretic Mobility-Shift Assays (EMSA) with nuclear protein extracts from NT2-D1 cells and a set of partially overlapping 100-150bp fragments indicate that three different fragments (N2, N5 and N7) within the 1kb PTE 1.2 element bind nuclear proteins in sequence specific fashion (Figure S10). Overall, it appears that the *CCND2* PTE is modular and depends on multiple sequence specific DNA binding proteins for PRC1 recruitment.

### PRC2 activity promotes PRC1 binding to the *CCND2* PTE

The PTE binds little PRC2 and H3K27me3 when *CCND2* is active. Considering that in TIG-3 cells, high levels of H3K27me3 and PRC2 are not sufficient to recruit much PRC1 to the region between the PTE and the *CCND2* TSS, we expected that at the low levels seen in NT2-D1 cells PRC2 and H3K27 methylation are dispensable for recruitment of PRC1 to the PTE. To our surprise, the CRISPR/Cas9-mediated knock-down of SUZ12 in NT2-D1 cells (Figure 7A, C) leads to a strong reduction of the PTE immunoprecipitation by the antibodies against PRC1 subunits (Figure 7B). Western-blot analysis argues that reduced immunoprecipitation of the *CCND2* PTE is not due to lower PRC1 abundance in the SUZ12 depleted cells (Figure 7A). In fact, NT2-D1 cells seem to require at least a low level of PRC2 for proliferation as after multiple attempts we failed to recover any cell lines completely deficient for SUZ12. In the clonal isolate used here (S1H6), the level of SUZ12 dropped to a few percent of that in the original NT2-D1 cells and the overall H3K27me3 is reduced about 10-fold (Figure 7A, C). Under these conditions SUZ12 and H3K27me3 are no longer detected at the *CCND2* PTE. At the same time, a very low SUZ12 ChIP signal is still detectable at the *ALX4* gene (used as a positive control example of a PcG-repressed gene in NT2-D1 cells). With this level of SUZ12 binding, *ALX4* retains significant immunoprecipitation with anti-H3K27me3 antibodies and ChIP signals for CBX2 and MEL18 are nearly unchanged (Figure 7B). Taken together, these results suggest that the physical presence of PRC2 or its enzymatic activity in some way contribute to robust recruitment of PRC1 to the *CCND2* PTE.

**Figure 7.**
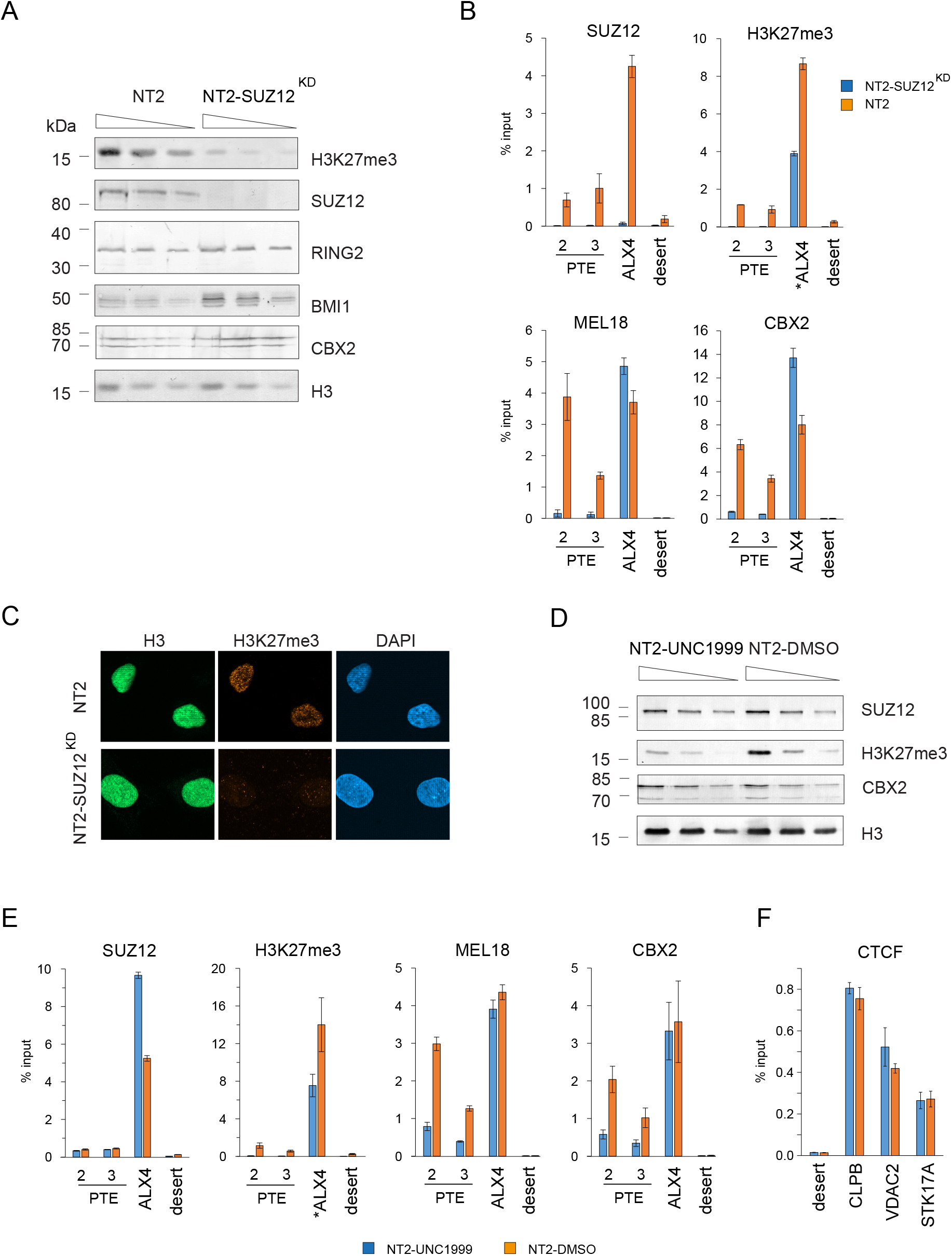
Catalytic activity of PRC2 promotes PRC1 binding to the PTE. **A.** Western-blot analysis of two-fold serial dilutions of nuclear protein from control and SUZ12 depleted cells shows that drastic reduction of SUZ12 and H3K27me3 does not affect the overall levels of PRC1 components. Here and in **D** detection of histone H3 is used as a loading control. **B.** ChIP analysis of SUZ12, H3K27me3, MEL18 and CBX2 binding at the *CCND2* PTE and control loci in SUZ12 depleted (blue bars) and control (red bars) cells. Positions of PTE amplicons are as indicated in Figure 2A. Here and in **E** and **F** histograms show the average and the scatter (whiskers) between two independent experiments. **C.** Immunofluorescent analysis of H3K27me3 reduction in SUZ12 depleted cells. Staining for total histone H3 is used as positive control and DAPI staining as nuclear marker. **D.** Western-blot analysis of two-fold serial dilutions of nuclear protein from UNC1999-treated and control cells shows that H3K27me3 is significantly reduced while the levels of SUZ12 and CBX2 remain unchanged. **E.** ChIP analysis of SUZ12, H3K27me3, MEL18 and CBX2 in control (red bars) and UNC1999-treated (blue bars) cells. **F.** ChIP analysis of CTCF occupancy at *CLPB*, *VDAC2*, *STK17A* genes and negative control locus (desert) in UNC1999-treated (blue bars) and control (red bars) cells.

To distinguish between the two possibilities, we inhibited PRC2 methyltransferase activity with a small molecule inhibitor UNC1999 shown to be highly specific for EZH2 and EZH1 and competitive with the cofactor SAM but not with the H3 substrate (61). Notably, UNC1999 does not lead to degradation of PRC2 and does not prevent its binding to chromatin (62). Consistent with observations in other cultured cell lines (61,62), twelve day treatment of NT2-D1 cells with recommended (2μM) concentration of UNC1999 leads to partial inhibition of the PRC2 activity without reducing overall levels of PRC2 and PRC1 (Figure 7D). Under these conditions, ChIP signals for H3K27me3 at the *CCND2* PTE drop to a background level but reduce less than two-fold at the control *ALX4* gene. At both sites the SUZ12 ChIP signals remain unchanged (Figure 7E). Importantly, following the UNC1999 treatment the MEL18 and CBX2 ChIP signals decrease 4-fold at the *CCND2* PTE but remain unchanged at *ALX4* (Figure 7E). To exclude the possibility that UNC1999 treatment causes indiscriminate loss of proteins bound next to active genes, we tested the binding of transcription factor CTCF upstream of the *CLBP*, *VDAC2*, *STK17A* genes. According to published expression array profile of the NT2-D1 cells (63), these genes are expressed within the same range (2 to 16 percent of *GAPDH*) as *CCND2*. As illustrated by Figure 7F, the CTCF ChIP signals are not affected by UNC1999 treatment. Taken together, our experiments suggest that enzymatic activity of PRC2 but not its physical presence are critical for robust binding of PRC1 to the PTE.

### PRC1 targeting elements may be common

The upstream region of the *CCND2* gene is organized in a particular way. It consists of the A/T-rich PTE, which is adjacent to the CpG-island that “links” the PTE to the *CCND2* TSS. How common is such “PTE – CpG-island – TSS” arrangement? The question is not trivial to address. First, we cannot unambiguously pinpoint other putative PTEs from genome-wide binding profiles since at transcriptionally inactive target genes PRC1 binding is rather broad. Second, it is possible that some of the PcG target genes use alternative ways to recruit PRC1. Taking this into account we reasoned that, if PTEs are sufficiently common and, like in the case of *CCND2*, correspond to sites of the strongest PRC1 binding, we should be able to detect the trace of the “PTE – CpG-island – TSS” arrangement by plotting aggregate distributions of di-nucleotide combinations around the highest BMI1/MEL18 binding peaks within PcG repressed genes.

Indeed, plotted such that all DNA sequences around the highest BMI1/MEL18 peaks on the human chromosomes 8, 11 and 12 are oriented to have the BMI1/MEL18 binding peak at the centre and the closest TSS to the right of the peak, the di-nucleotide distributions show three striking features (Figure 8A). First, the DNA sequences underneath the BMI1/MEL18 peaks tend to be AA-rich but CG-poor. Second, the DNA around BMI1/MEL18 peaks is CG-rich with clear bias towards TSSs. Third, the DNA encompassing BMI1/MEL18 peaks is depleted of TG/CA nucleotides. This is in contrast to average di-nucleotide distributions in upstream regions of human genes (Figure S11). These control distributions show no AA-rich hot-spots and matching CG-poor gap. They also display no TG/CA depletion. Regions upstream of TSS appear generally CG-rich, however their CG enrichment is lower than that downstream of the BMI1/MEL18 peaks. The picture of di-nucleotide distributions around BMI1/MEL18 peaks parallels the arrangement seen at the *CCND2* locus (compare Figure 8A to Figure 1C). This suggests that many of the DNA sequences underneath MEL18/BMI1 peaks contain AA-rich elements similar to the *CCND2* PTE and that the arrangement “AA-rich MEL18/BMI1 peak – CpG-island – TSS” is common for many PcG target genes.

**Figure 8.**
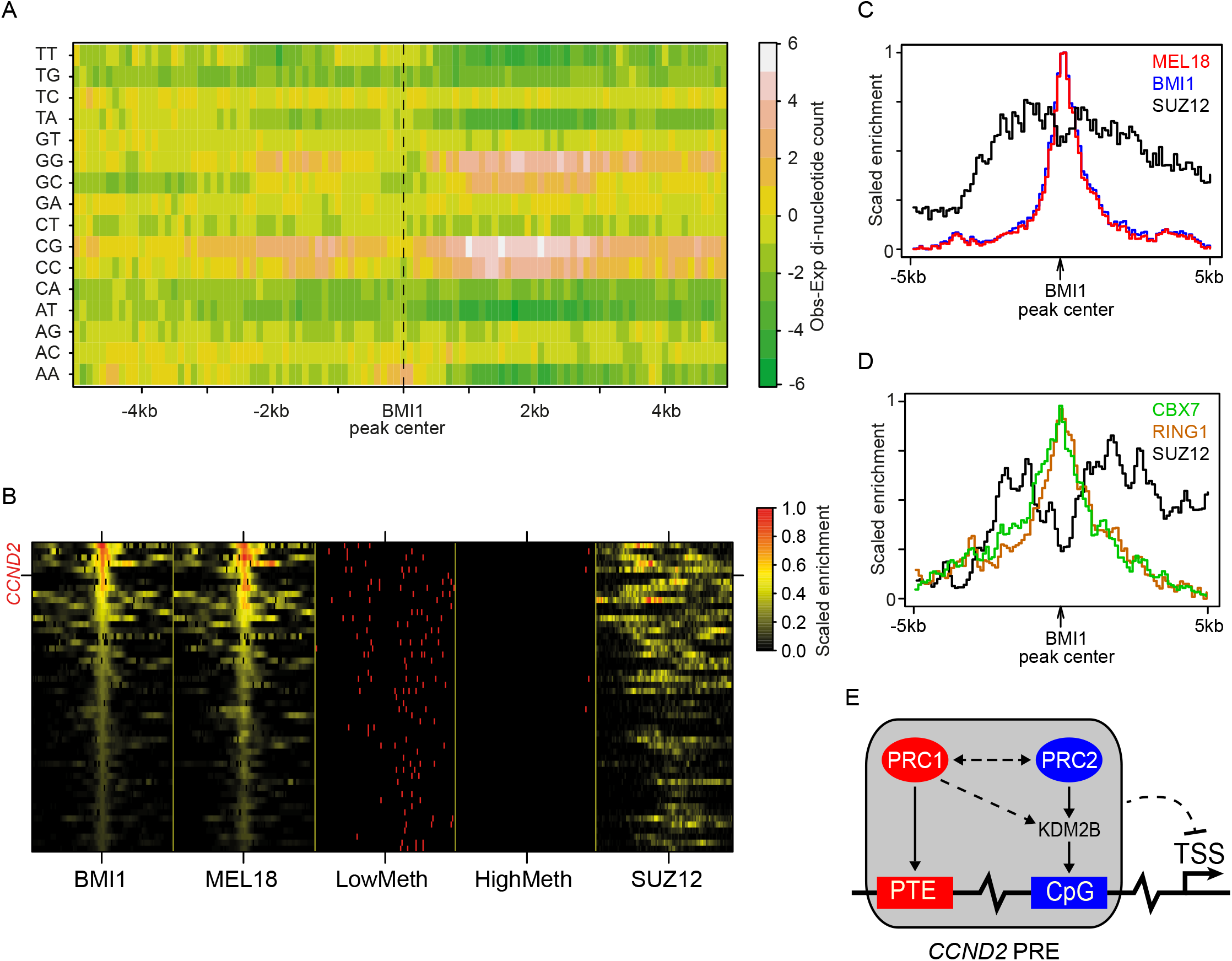
Combined presence of PTE and CpG-islands paralleled by segregation of PRC1 and PRC2 binding are common features of PcG target genes. **A.** The heat-map of di-nucleotide distributions around high confidence BMI1 peaks. Each square represents the difference between the mean di-nucleotide count within a 100 bp window (observed) and the average count throughout the genome (expected). **B.** Heat-maps showing the enrichment of BMI1, MEL18, the methylation status of the CpG-islands and the enrichment of SUZ12 around BMI1 peaks. Here and in **C** and **D** the values are adjusted by subtracting the genomic background and linear scaling between 0 and 1. **C.** The average ChIP signals for MEL18 (red), BMI1 (blue) and SUZ12 (black) in NT2-D1 cells are plotted in 100bp increments around high confidence BMI1 binding sites. Note the clear difference between the BMI1/MEL18 and the SUZ12 profiles. **D.** The relative shift between PRC1 components and SUZ12 is also seen when the average ChIP signals for CBX7 (green), RING1 (orange) in primary fibroblasts and the SUZ12 signals (black) from embryonic stem cells are plotted around the same BMI1 binding positions. **E.** A speculative model of *CCND2* regulation by the PcG proteins. Our results suggest that *CCND2* is equipped with two DNA elements: the PTE and the CpG-island, which serves as focus of KDM2B and PRC2 binding. When *CCND2* is transcriptionally inactive, CpG-anchored KDM2B may acquire RING2-PCGF1 dimer with the help of PTE-anchored PRC1. We propose that the PTE and the CpG-island cooperate to act as the *CCND2* PRE.

PRC2 binding was linked to unmethylated CpG islands (41–43,64) and in TIG-3 cells, where the *CCND2* gene is repressed, the ChIP signal for SUZ12 is offset from the PTE to the adjacent CpG-rich region. Strikingly, the average SUZ12 distribution around MEL18/BMI1 peaks in NT2-D1 cells (52) parallels that of the unmethylated CpG di-nucleotides (Figure 8B). We see that the SUZ12 binding is generally offset to the sides of the MEL18/BMI1 peaks (Figure 8C). This observation is not unique for NT2-D1 cells. Comparing the ChIP-seq profiles of CBX7 and RING1 in primary fibroblasts (65) to the ChIP-seq profile of SUZ12 in human embryonic stem cells (52) we see essentially the same picture (Figure 8D). Overall, these results suggest that a significant fraction of human PcG target genes recruit PRC complexes via a combination of AA-rich PTEs and CpG-islands.

To further evaluate the hypothesis that *CCND2* is not the only gene equipped with a PTE, we searched for more examples. As demonstrated by the *CCND2* case, a PTE can be pinpointed as localized PRC1 binding site when its target gene is transcriptionally active and has little PRC2 and H3K27me3. We expect such cases to be rare. First, a large fraction of Polycomb target genes encodes tissue specific transcription factors (36,50,66,67) and, therefore, most of these genes will be inactive in any given cell type. Second, there is likely more than one way to target PRC1 and, therefore, not all Polycomb target genes are equipped with PTEs. To screen for additional PTE-equipped genes, we performed a rapid mapping of MEL18 and SUZ12 in NT2-D1 cells genome-wide with ChIP-seq. From these data, we identified MEL18-bound regions that had an annotated TSS within 20kb distance and displayed MEL18 ChIP-seq signal comparable to that at *CCND2*. These regions were further ranked by MEL18/SUZ12 ChIP-seq signal ratios. Consistent with published ChIP-chip profiles used to pinpoint *CCND2* PTE, the *CCND2* locus was ranked second and had the highest rank from all loci on chromosomes 8, 11 and 12 that were tiled by our Affymetrix array (Table S1). To be conservative in interpreting our non-extensive ChIP-seq profiles, we focused on the site that scored better than *CCND2* (i.e. was ranked first and had higher MEL18 ChIP-seq signal). It corresponded to a narrow MEL18 bound region approximately 10kb downstream of the *ZIC2* gene (Figure S12A). Like the *CCND2* PTE, this putative PTE region is CpG-poor but surrounded by CpG-islands. RT-qPCR analysis confirmed our expectation that *ZIC2*, the likely target of PcG regulation, is highly expressed in NT2-D1 cells. Fortunately, *ZIC2* is inactive in TIG-3 cells providing essential comparison (Figure S12B). The putative *ZIC2* PTE is traversed by a long non-coding RNA *LINC00554* of unknown function, which is not detectable in either cell line (Figure S12B).

Detailed ChIP-qPCR analysis shows that, just like *CCND2*, the *ZIC2* locus binds PRC1 at the putative PTE in both NT2-D1 and TIG-3 cells regardless of transcriptional status of the *ZIC2* gene (Figure S12C). However, the antibodies against SUZ12 and H3K27me3 efficiently immunoprecipitate the locus only from the chromatin of TIG-3 cells, where *ZIC2* is transcriptionally inactive (Figure S12C). In line with what is seen at the *CCND2* gene (Figure 2B) and at an average PcG target locus (Figures 8C-D), the SUZ12 and H3K27me3 ChIP signals in TIG-3 cells are offset from the putative PTE into neighboring CpG-rich regions. Taken together, the *ZIC2* example and the bioinformatic analyses above strongly suggest that *CCND2* is not the only human gene equipped with a PRC1 targeting element.

## Discussion

Epigenetic repression by Polycomb Group mechanisms is critical for development of all multicellular animals and is frequently disrupted in cancers (4,68,69). Yet, how it is targeted to specific genes is not well understood. More is known about the process in *Drosophila*. There we know that the targeting is mediated by compact DNA elements (PREs), that PREs are sufficient to recruit all Polycomb complexes and that the recruitment requires combinatorial action of sequence specific DNA binding adaptor proteins (32,70,71). We also know that PREs can bind PRC1 independently of PRC2 and that, at many PREs, PRC1 has to be present for robust PRC2 binding (38). Targeting PcG repression in mammalian cells is likely somewhat different. First, mammalian genomes encode greater variety of RING complexes that contribute to PcG repression. Second, as a consequence of CpG methylation, mammalian genomes are AT-rich and their genes and corresponding regulatory elements are often embedded within CpG-rich islands protected from DNA methylation. This feature, absent from *Drosophila*, seems to help targeting of mammalian PRC2 complexes (41–43,64). On the other hand, many of the same developmental genes are regulated by Polycomb mechanisms in flies and mammals. This suggests that some of the common target genes have been regulated by PcG complexes in the last common ancestor of vertebrates and insects and that at least part of the original targeting mechanism is still in place in both animal branches.

Here we used the peculiar binding of PRC1 to the transcriptionally active *CCND2* gene as an entry point to look for similarities and differences in the targeting of PcG complexes in *Drosophila* and humans. We show that, similar to flies, PRC1 complexes are autonomously targeted to *CCND2* by a compact DNA element (PTE). Like fly PREs, the *CCND2* PTE consists of several partially redundant modules recognized by distinct sequence specific DNA binding activities. However, the *CCND2* PTE also differs from *Drosophila* PREs. PREs can robustly recruit both PRC1 and PRC2 and therefore are self-sufficient to install PcG repression. In contrast, the *CCND2* PTE seems to recruit PRC1 very well but PRC2 only poorly. Comparison of PRC1 and PRC2 binding to *CCND2* in alternative transcriptional states suggests that robust targeting of both complexes involves multiple DNA elements: A/T-rich PTE, responsible for PRC1 recruitment, and adjacent CpG-island that acts as focus of PRC2 targeting. We propose that the combination of these two elements corresponds to a functional analogue of a *Drosophila* PRE (Figure 8E). In this view, the targeting of PRC1 by a compact DNA element is a common ancestral feature while the targeting of PRC2 has diverged. Flies, which lost DNA methylation, must have evolved different means to target PRC2 in a way that is more hardwired to the targeting of PRC1 (38,72).

Consistent with this view, unmethylated CpG-islands are recognized by the mammalian RING2-PCGF1-KDM2B complex via the CXXC domain of KDM2B (24,25,51). This complex mono-ubiquitylates H2AK119 and ubiquitylation is directly recognized by PRC2 and may help to recruit it (23,29,48). In contrast, the fly genome encodes no PCGF1 protein and the corresponding KDM2B complex does not exist. Furthermore, the fly H2AK118ub is not sufficient and, at least at some genes, not necessary for stable PRC2 binding or Polycomb repression (12,14,38).

In TIG-3 cells immunoprecipitation of the *CCND2* CpG-island with H2AK119ub antibodies correlates with strong ChIP signals for PRC2 and H3K27me3. However, comparable H2AK119ub signal over the PTE in NT2-D1 cells does not. This suggests that H2AK119ub is not sufficient to recruit much PRC2 or that it is prevented from doing so by a process linked to the *CCND2* expression. Interestingly, KDM2B seems to bind the *CCND2* CpG-island regardless of the transcriptional status of the gene. However, it is accompanied by RING2 only when *CCND2* is repressed. It is tempting to speculate that, when *CCND2* is transcriptionally inactive, the CpG-anchored KDM2B “receives” its RING2 from PRC1 anchored at the nearby PTE to form RING2-PCGF1-KDM2B ubiquitylase.

In NT2-D1 cells the endogenous and transgenic *CCND2* PTE binds little PRC2 and displays little H3K27 tri-methylation. Paradoxically, the binding of PRC1 to the PTE is strongly reduced when PRC2 enzymatic activity is impaired. This dependence is hard to explain by postulating that H3K27me3 serves as a platform for PRC1 binding and therefore the binding is abolished when PRC2 activity and H3K27me3 are lost. First, the PRC1 ChIP signals at the PTE are very similar between NT2-D1 and TIG-3 cells although the extent of immunoprecipitation with anti-H3K27me3 antibodies is very different. More importantly, in TIG-3 cells a much higher level of H3K27me3 over the CpG-island does not lead to PRC1 binding comparable to that at the PTE suggesting that H3K27me3 is not sufficient for robust recruitment. Conceivably, H3K27me3 serves a “catalytic” function by binding chromodomain of a CBX subunit and introducing allosteric changes to the PRC1 complexes that promote interaction with DNA binding adapter proteins. Alternatively, the effect may be indirect and stem not from the PRC2 complex bound at the PTE but from the loss of untargeted PRC2 activity. This activity was shown to globally suppress chromatin accessibility and could be, in some way, beneficial for PRC1 binding (13,14). The puzzling PRC1 dependence on PRC2 for binding the *CCND2* PTE is not without a precedent. A DNA element upstream of the mouse *MafB* gene, which resembles the *CCND2* PTE in its ability to recruit PRC1 but not PRC2, was also reported to inexplicably lose PRC1 binding after the SUZ12 knock-down by RNAi (45).

Serendipitous discovery of the PTE turned the *CCND2* locus into a fruitful model that gave us mechanistic insights in principles of PcG protein targeting and shed light on the parallels between the processes in *Drosophila* and mammals. We have little doubt that *CCND2* has much more to offer. Of obvious interest is the nature of the DNA binding proteins implicated at the *CCND2* PTE as those are likely to be involved in the recruitment of PRC1 complexes to other human genes. Other questions that could be addressed using *CCND2* as a model include the roles of H2AK119 ubiquitylation and the RING2-PCGF1-KDM2B complexes in targeting of PRC2 and the identity of the processes that prevent PRC2 binding when a target gene is active. It will also be interesting to explore whether the *CCND2* PTE, or PTEs in general, have preferences for PRC1 variants with different CBX subunits. For example, CBX2 has the chromodomain coupled to an AT-hook and in early mouse embryos the two domains cooperate to target CBX2 to paternal AT-rich satellite DNA (73). Conceivably, the same feature may help the CBX2-containing PRC1 complexes to bind the AT-rich PTEs. Finally, *CCND2* is an oncogene (74–76) and multiple lines of evidence link PcG mechanisms to cancer progression (69,77). It is still puzzling why some tumour types depend on the overexpression of PcG proteins while others require the loss of PcG function. The oncogenic effect of the overexpression has to some extent been explained by erroneous silencing of the *INK4A/ARF* locus (78) but the link between malignant transformation and the loss of PcG proteins remains elusive. We speculate that a failure to repress *CCND2* may, at least in part, explain this link. From this we predict that the *CCND2* PTE may be disrupted in some cancers that overexpress *CCND2* but otherwise have the PcG function intact.

## Materials and Methods

### Transgenic constructs

Lentiviral constructs were produced by *in vitro* recombination of fragments of interest into the Eco47III site of the pLenti-CMVTRE3G-eGFP-ICR-Puro or pLenti-ICR-Puro vectors. *In vitro* recombination was done using In-Fusion HD system (Clontech). All fragments were amplified using high-fidelity Pfu DNA polymerase (Thermo Scientific) and human or mouse genomic DNA or, in case of *CCND2* PTE sub-fragments, DNA of the PTE 2.4kb construct as a template. PCR primers and their sequences are indicated in Tables S2, S3.

Plenti-CMVTRE3G-eGFP-ICR-Puro was generated based on the pLenti-CMVTRE3G-eGFP-Puro backbone (79). As first step, pLenti-CMVTRE3G-eGFP-Puro was cut with HpaI and EcoRI and recombined with two DNA fragments produced by PCR with following pairs of primers: 5’-TCGACGGTATCGGTTAACTT-3’; 5’-AGCGCTAGTCTCGTGATCGATAAA-3’ and 5’-CACGAGACTAGCGCTGAGAGTTGG-3’; 5’-CTACCCGGTAGAATTCCACGTGGGGAG-3’ and DNA of pLenti-CMVTRE3G-eGFP-Puro as a template. This step introduced unique PmlI site between *eGFP* and *Pac* (puromycin resistance gene) and unique Eco47III site upstream of the *eGFP* gene. As second step, the resulting construct was digested with PmlI and recombined with the mouse *Igf2/H19* ICR insulator sequence PCR amplified from mouse genomic DNA using the forward 5’-CCGGTAGAATTCCACTGTCACAGCGGACCCCAACCTATG-3’ and the reverse: 5’-GGGCCGCCTCCCCACTCGTGGACTCGGACTCCCAAATCA-3’ primers. The native *Igf2/H19* ICR sequence contained one Eco47III site. As final step, this site was removed by cutting the above construct with Eco47III and recombining it with the DNA fragment produced by PCR with the following primer pair: 5’-GATCACGAGACTAGCGCTGAGAG-3’ and reverse 5’-TTTTCACACAATGGCGCTGATGGCC-3’, using DNA of the above construct as a template.

We originally planned to use the pLenti-CMVTRE3G-eGFP-ICR-Puro based constructs and integrate them into the NT2-D1 cells that had been modified to express TetR protein from constitutive CMV promoter. In theory this should have allowed us to induce eGFP by adding the doxycycline to the media. Unfortunately, we have soon discovered that the CMV promoter is not active in NT2-D1 cells. Therefore to simplify and reduce the size of the transgenes we have removed the TRE3G-eGFP part from pLenti-CMVTRE3G-eGFP-ICR-Puro to yield the pLenti-ICR-Puro vector (Figure S6A). This was done by digesting pLenti-CMVTRE3G-eGFP-ICR-Puro with EcoRV and Eco47III and recombining it with short dsDNA fragment produced by the annealing of the 5’-GATCACGAGACTAGCGCTGAGAGTTGGCTTCACGTGCTAGACCCAGCTTTC-3’ and 5’-GAAAGCTGGGTCTAGCACGTGAAGCCAACTCTCAGCGCTAGTCTCGTGATC-3’ oligonucleotides. Side-by-side comparison of the PRC1 binding to transgenic PTE 1.2 in either vector showed that the presence of the CMVTRE3G-eGFP part had no effect on the recruitment of PRC1. The analyses on Figure 3B-C and Figure S6B were done with transgenic constructs based on pLenti-CMVTRE3G-eGFP-ICR-Puro. All other analyses used transgenes based on pLenti-ICR-Puro.

### Cell culture and lentiviral transduction

NTERA-2 (NT2-D1 ATCC^®^ CRL-1973^TM^), TIG-3 (gift from Dr. K. Helin, BRIC), 293T (ATCC^®^ CRL-3216^TM^) and F9 (ATCC^®^ CRL-1720^TM^) cells were cultured in high-glucose DMEM (Gibco) supplemented with 10% Fetal Bovine Serum (Sigma), Penicillin/Streptomycin (Gibco) and 1.5g/L Sodium Bicarbonate (Sigma) at 37°C in atmosphere of 5% CO_2_. The F9 cells were plated in T-flasks pre-treated with 0.1% gelatin in H_2_O for 1 hour prior to plating.

To produce viral particles, 5×10^6^ 293T cells were plated in a 75cm^2^ flask 24 hours prior to transfection. The packaging plasmids pCMV-dR8.2dvpr (6μg) and pCMV-VSV-G (3μg) (gift from of M. Roth, UMDNJ) were combined with a transfer construct (9μg) and cotransfected using X-tremeGene HP (Roche) at 1:3 ratio of DNA to transfection reagent. After 24 hours of incubation, the medium was changed. Lentiviral supernatant was collected after another 24 hours, filtered (0.45 μm filters) and used for infection directly or stored at −80°C.

For lentiviral infection, cells were plated at a confluence of 40 to 60% 24 hours in advance. Viral supernatant was added in serial dilution to cells in combination with 8μg/ml Polybrene (Millipore). After overnight incubation, the medium was changed to remove Polybrene. Transduced cells were selected for 14 days by growth on culture medium supplemented with 4μg/ml puromycin (Invitrogen).

To generate the SUZ12 knock-out NT2 cell line the SUZ12g1.1 (caccgGGTGGCGGCGGCGACGGCTT) and SUZ12g1.2 (aaacAAGCCGTCGCCGCCGCCACCc) DNA oligonucleotides were annealed and cloned into lentiCRISPRv2 plasmid (Addgene #52961) linearized by digestion with *BsmBI*. The construct was introduced in NT2-D1 cells by lentiviral infection and transduced cells selected by growth on the culture medium supplemented with 4μg/ml puromycin (Invitrogen). Cells were cloned and knock-down assayed by Western blot and immunofluorescence with the antibodies listed in Table S4.

To inhibit EZH2/EZH1, the NT2-D1 cells were grown in complete DMEM supplemented with 2μM of UNC1999 (Cayman Chemical Company #14621) or 0.2% DMSO as a negative control. The media were replaced every second day for 12 days and cells crosslinked for ChIP analyses. Fractions of the same cell cultures were taken before crosslinking and their total nuclear protein analyzed by western blot.

### ChIP and RT-qPCR analyses

ChIP reactions were performed as described in (36,49). For some experiments the procedure was modified to include additional crosslinking with EGS (ethylene glycol bis(succinimidyl succinate)). Cells were scraped, resuspended in 10ml of 1xPBS and EGS solution in DMSO added to final concentration of 2mM. The crosslinking reaction was allowed to proceed for 1 hour at room temperature after which formaldehyde was added to final concentration 1% and crosslinking continued for another 15 minutes at room temperature. The reaction was stopped by adding an excess of glycine. The antibodies used for ChIP are listed in the Table S4. Total RNA from cultured cells was isolated using TRI Reagent (Sigma) according to manufactures instructions. cDNA was prepared with the First Strand cDNA synthesis kit (Thermo Scientific) using 2μg of RNA and random hexamer primers and purified as described (37). qPCR analysis of cDNA and ChIP products was performed essentially as described in (37,80) except that iQ5 Real-Time PCR Detection system (Bio-Rad) and KAPPA SYBR FAST qPCR Kit (Kappa Biosystems) were used for all analyses. The primers used for qPCR analyses are described in Tables S5, S6, S7. Sequencing of ChIP products was done as described in (38) except that NEBNext Ultra II DNA Library Prep Kit for Illumina (NEB #E7645S) was used for sequencing library preparation.

### Electrophoretic Mobility-Shift Assay

To prepare the nuclear extract, the NT2-D1 cells from four 182.5cm^2^ T-flasks were disrupted by resuspension in 400μl of hypotonic Cell Lysis Buffer (5% Sucrose, 5mM Tris-HCl pH8, 5mM NaCl, 1.5mM MgCl_2_, 0.1% Triton X-100, 1mM PMSF, 2mM dithiolthreitol, 1x Protease Inhibitor Cocktail). After discarding the cytosolic fraction, nuclei were extracted for 1 hour at 4°C with 110μl of Nuclear Extract Buffer (20mM HEPES, 20% Glycerol, 500mM NaCl, 1.5mM MgCl_2_, 0.2mM EDTA, 0.5mM dithiolthreitol, 0.5mM PMSF, 1x Protease Inhibitor Cocktail). The nuclear extract was cleared by centrifugation for 20 minutes at 16000g, 4°C and used immediately or stored at −80 °C.

dsDNA fragments were labeled with αP^32^ dATP (Perkin Elmer) using PCR and the DNA of the PTE 2.4kb construct as a template and purified by passing through Illustra MicroSpin S300 HR Columns (GE Healthcare). The corresponding PCR primers are indicated in the Table S8.

Binding reactions (final volume of 20μl) were assembled by combining 5μl of 4x binding buffer (80mM HEPES pH7.4-7.9, 20mM MgCl_2_, 20% Glycerol, 400mM NaCl, 4mM dithiolthreitol, 4mM EDTA), 5μg to 10μg of nuclear protein, 1μg to 5μg of Salmon Sperm DNA (Life Technologies), 40fmol of labeled probe and 150 fold excess of specific or nonspecific competitor. The binding was allowed to proceed for 20 minutes at room temperature after which samples were run on 8% acrylamide gel (29:1 acrylamide to bis-acrylamide solution) on 1xTBE for 4 to 6 hours at 160 V. The resulted gels were dried for 2 hours and exposed to x-ray film (AGFA Healthcare).

### Computational analyses

#### Definition of bound regions

The MEL18 and BMI1 bound regions were defined as coordinates of clusters of microarray features that satisfied the following three criteria. i) smoothed ChIP/Input hybridization intensity ratios of the features were above the 99.8 percentile cut-off; ii) the maximum distance to the neighboring feature above the intensity cut-off was equal or greater than 200bp; and iii) the length of the cluster was equal or greater than 200bp. The peak center of a bound region was set at the center of the 5 consecutive microarray features with the highest hybridization intensity values. Regions of 1kb, centered on overlapping binding peaks of MEL18 and BMI1 were considered as high-confidence MEL18/BMI1 binding sites. To avoid overlapping regions in the analysis for figures 1C, 8A and 8B we increased the ii-criterion above to 20kb. CpG-islands were defined using default parameters in the EMBOSS package (81).

#### Motif analysis

The 317 most predictive words from the 1kb MEL18/BMI1 high-confidence regions derived by multivariate modelling as described in (59,60) were pair-wise aligned and each alignment was assigned a score reflecting the maximum number of identical nucleotides in the alignment. Based on these scores we generated a hierarchical tree (Euclidian distance and complete linkage) using Cluster 3.0 (82). The tree was divided into eight groups and the words from each group were realigned using Muscle (83). The aligned words were used to build the position weight matrix (PWM) following the method proposed by Staden (84). As final step, the resulted PWMs were optimized with the Bound/Surveyed Sequence Discrimination Algorithm (85). The prediction of potential binding sites for the YY1, RUNX1/CBFβ, REST and SNAIL proteins was done as described in (86).

#### ChIP-sequencing data analysis

Sequencing reads were aligned to the Hg19 human reference genome with bowtie2 (87) using default parameters. Resulted SAM files were filtered using samtools (88) to remove reads with q-scores lower than 10. The reads were extended to 180bp (the average insert size in sequencing libraries) and read density profiles generated using Pyicos (89). To compare MEL18 and SUZ12 binding patterns, the read density profiles were filtered to remove all data points below 99^th^ percentile and bound regions defined as clusters of five or more genomic positions separated by less than 501bp that had read densities greater than 0. For each detected region, the “binding” value was set to an average of the five consecutive highest read density values and the position of the corresponding binding peak to their middle coordinate. To calculate MEL18/SUZ12 binding ratios, the binding value of each MEL18 peak was divided by that of the “strongest” SUZ12 binding peak in the 10kb neighborhood.

#### External data

The ChIP-chip profiles of MEL18, BMI1, EZH2 and H3K27me3 on Chromosomes 8, 11 and 12 of NT2-D1 cells were from GSE41854 (49). The NT2-D1 DNA methylation profiles were from GSM683929 and GSM683804. The SUZ12 ChIP-seq profiles in NT2-D1 and human embryonic stem cells were from GSM935423 and GSM1003573, respectively. The CBX7 and RING1 ChIP-seq profiles in primary human fibroblasts were from GSE40740.

## Data access

The ChIP-seq data are available from GEO at the accession number GSE101538

## Acknowledgements

We are grateful to Dr. Kristian Helin for the gift of TIG-3 cells and Dr. Monica Roth for the gift of lentiviral packaging plasmids. We thank Dr. Haruhiko Koseki and Dr. Robert Klose for generous gifts of anti-RING2 and anti-KDM2B antibodies. We are grateful to Dr. Jana Šmigová for evaluation of anti-CTCF antibodies. This work was supported in part by grants from Åke Wiberg and Magnus Bergvall foundations to PS, grants from Cancerfonden, Umeå Universitet Insamlingsstiftelsen and Strategic Research grant from Medical Faculty, Umeå University to YBS and grants from Knut and Alice Wallenberg Foundation and Kempestiftelserna to EpiCoN (YBS and PS co-PIs).

## Author contributions

S.C. generated all transgenic cell lines and performed ChIP analyses of human transgenic lines and EMSA. S.C. and J.I.B. performed ChIP analysis of mouse *Ccnd2* PTE. S.C. and T.G.K. analyzed PcG protein binding to *CCND2* locus in NT2-D1 and TIG-3 cells. T.G.K. performed PRC2 knock-down experiments, genome wide mapping of MEL18 and SUZ12 and characterized *ZIC2* PTE. J.I.B. helped with characterization of the PRC2 knock-down cell lines. S.N. performed most of the computational analyses. P.S. supervised S.N. and performed computational analyses. S.C., S.N., T.G.K., J.I.B., P.S. and Y.B.S. interpreted the data. Y.B.S. conceived and supervised the project and wrote the manuscript with input from all co-authors.

## Supplementary figure legends

**Figure S1.**
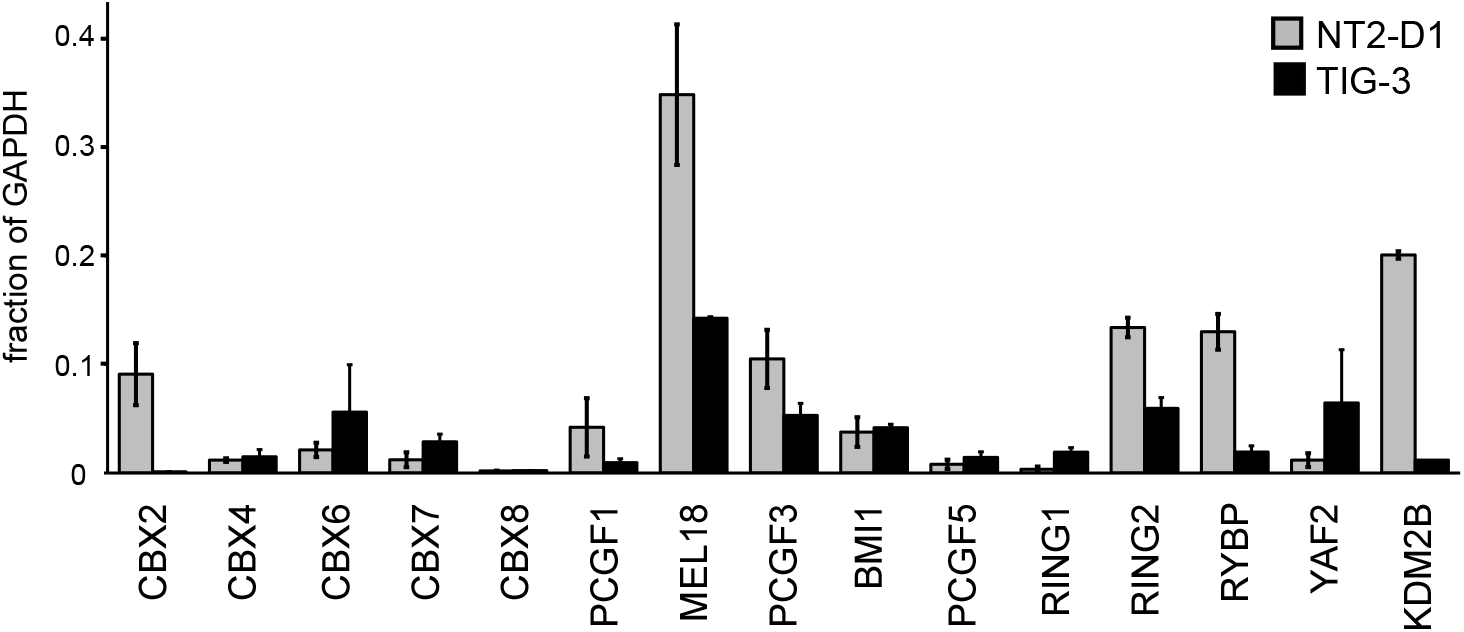
The mRNA levels for genes encoding components of various RING1/RING2 complexes. Histograms show the average and the scatter (whiskers) between two independent RT-qPCR experiments. Values are normalized to the expression of the housekeeping *GAPDH* gene.

**Figure S2.**
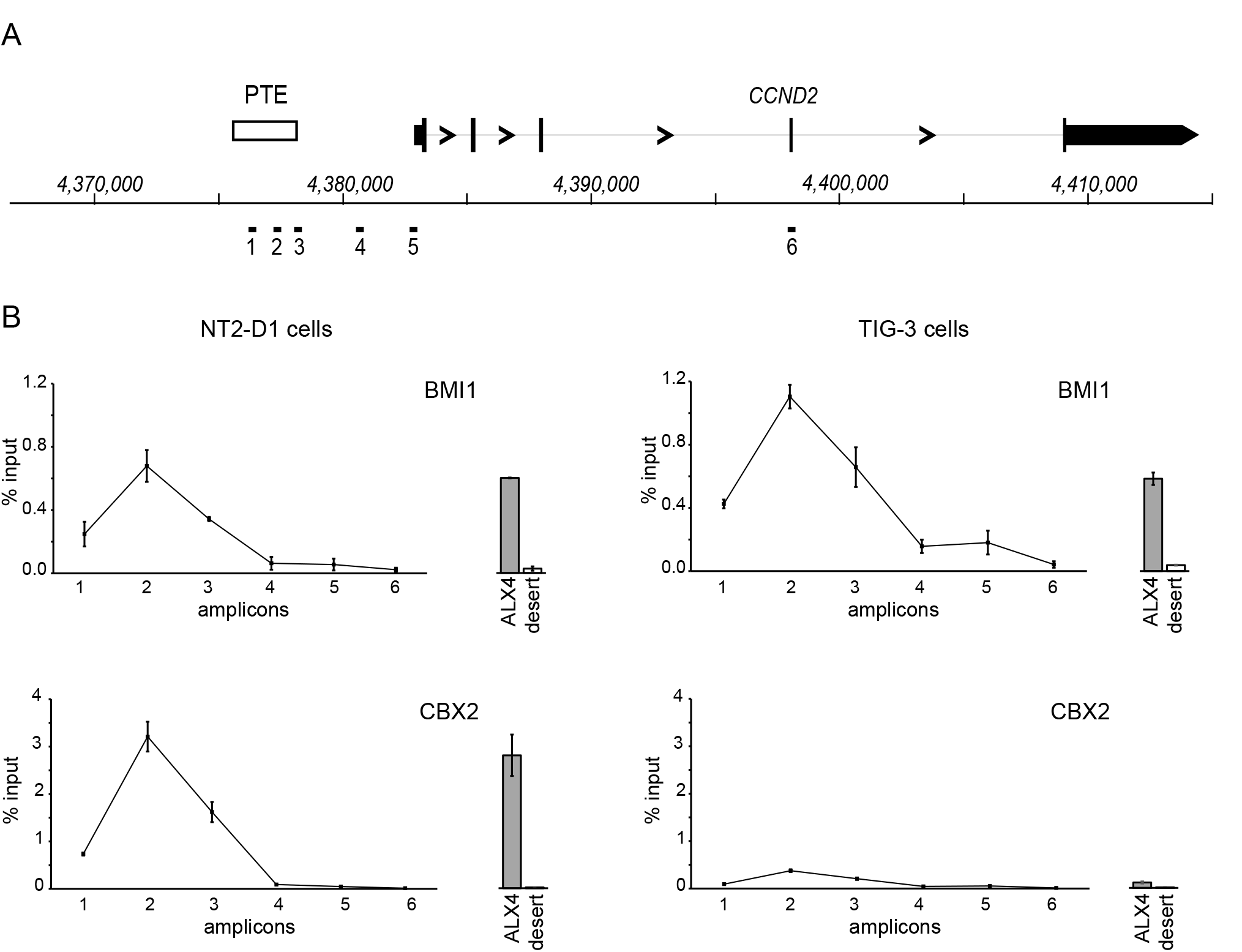
BMI1 and CBX2 at the active and repressed *CCND2* gene. **A.** The schematic of the *CCND2* locus. Positions of PCR amplicons analyzed in **B** are shown below the coordinate scale. **B.** ChIP profiles of BMI1 and CBX2 in NT2-D1 (left column) and TIG-3 (right column) cells. The immunoprecipitation of the *ALX4* gene and a gene desert region were used as positive and negative controls. All histograms and graphs show the average and the scatter (whiskers) between two independent experiments. The CBX2 ChIP signals anywhere in the *CCND2* locus or in the positive control locus (*ALX4*) in TIG-3 cells are very low likely because the corresponding gene is poorly expressed in these cells.

**Figure S3.**
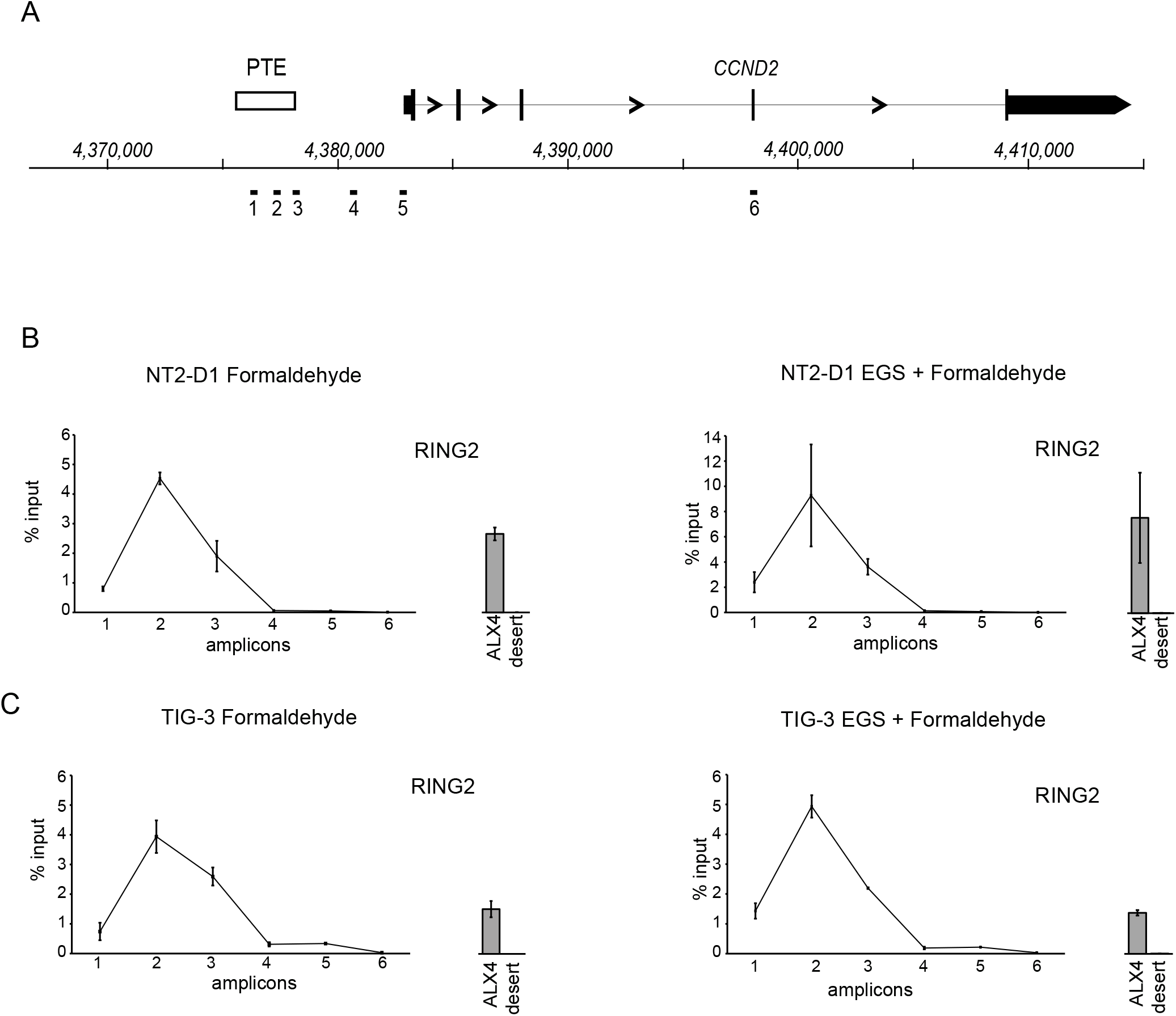
Immunoprecipitations of RING2 from chromatins crosslinked with formaldehyde or with a combination of EGS and formaldehyde give the same result. **A.** The schematic of the *CCND2* locus. RING2 ChIP profiles from variously crosslinked NT2-D1 (**B.**) and TIG-3 (**C.**) cells. The immunoprecipitation of the *ALX4* gene and a gene desert region were used as positive and negative controls. All histograms and graphs show the average and the scatter (whiskers) between two independent experiments.

**Figure S4.**
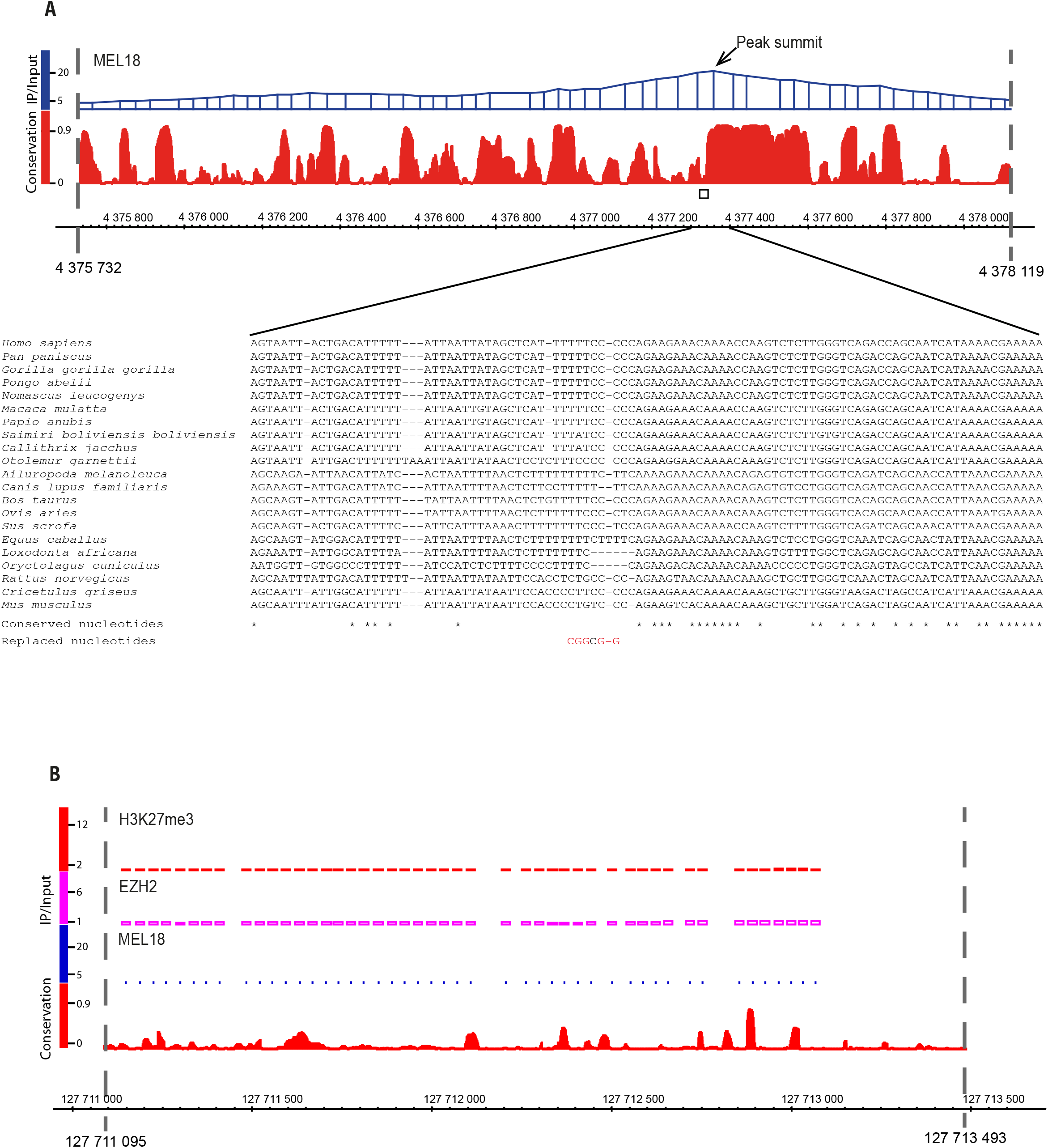
Evolutionary conservation of the DNA sequence underneath the *CCND2* MEL18/BMI1 peak. **A.** The comparison of the MEL18 binding profile with the PhastCons sequence conservation scores based on the alignment of 46 mammalian genomes highlights a small stretch of nucleotides close to the summit of the MEL18 peak that shows little conservation (rectangular box). The lower panel shows the location of five nucleotides (shown in red under the sequence alignment) substituted to generate an annealing site for a transgene-specific PCR primer. The dashed lines mark the boundaries of the 2.4kb DNA fragment used in transgenic assays. The genomic coordinates are in release Hg 19. **B.** The gene desert region from Chromosome 12 selected as negative control. The dashed lines mark the boundaries of the DNA fragment used in transgenic assays. The ChIP-chip signals for EZH2, H3K27me3 and Mel18 indicate the lack of binding paralleled by poor evolutionary conservation.

**Figure S5.**
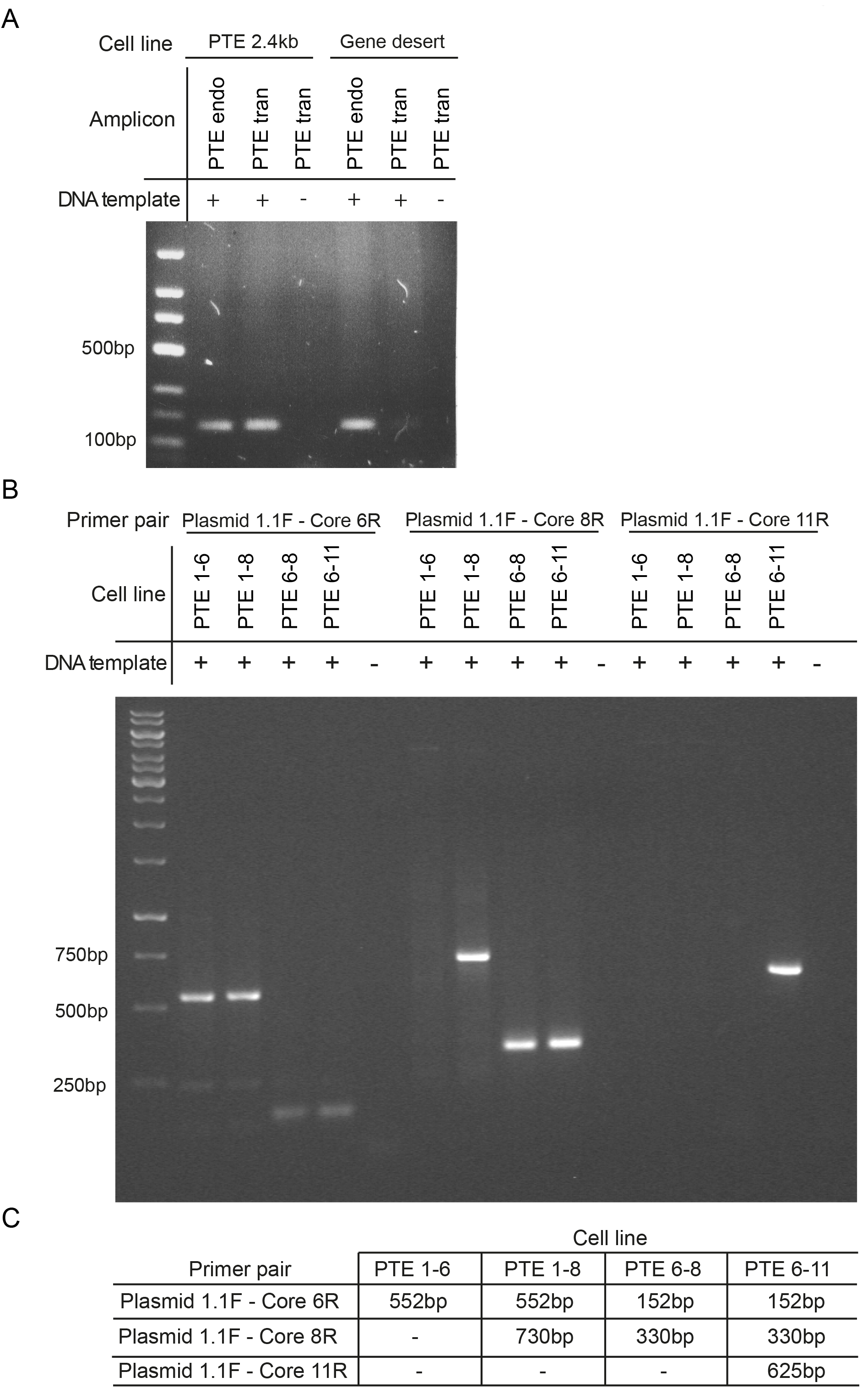
PCR genotyping of transgenic cell lines. All transgenic lines were PCR genotyped using ChIP input material as a DNA template. **A.** The results of a representative analysis of PCR products shows that transgene-specific primers (PTE tran) amplify the product of expected size from genomic DNA of the cells transduced with PTE 2.4kb construct but not with the control (Gene desert) construct. This verifies both the identity of the cell lines and the specificity of the “PTE tran” primers. **B.** The analyses similar to those in **A.** were performed for cell lines transduced with PTE sub-fragments. Sequences of corresponding primers are indicated in the Table S3. **C.** Anticipated results of PCR genotyping are summarized in the table that indicates expected sizes of PRC products. The dash indicates that no product should be produced. Note that the results in **B.** match the expected outcomes.

**Figure S6.**
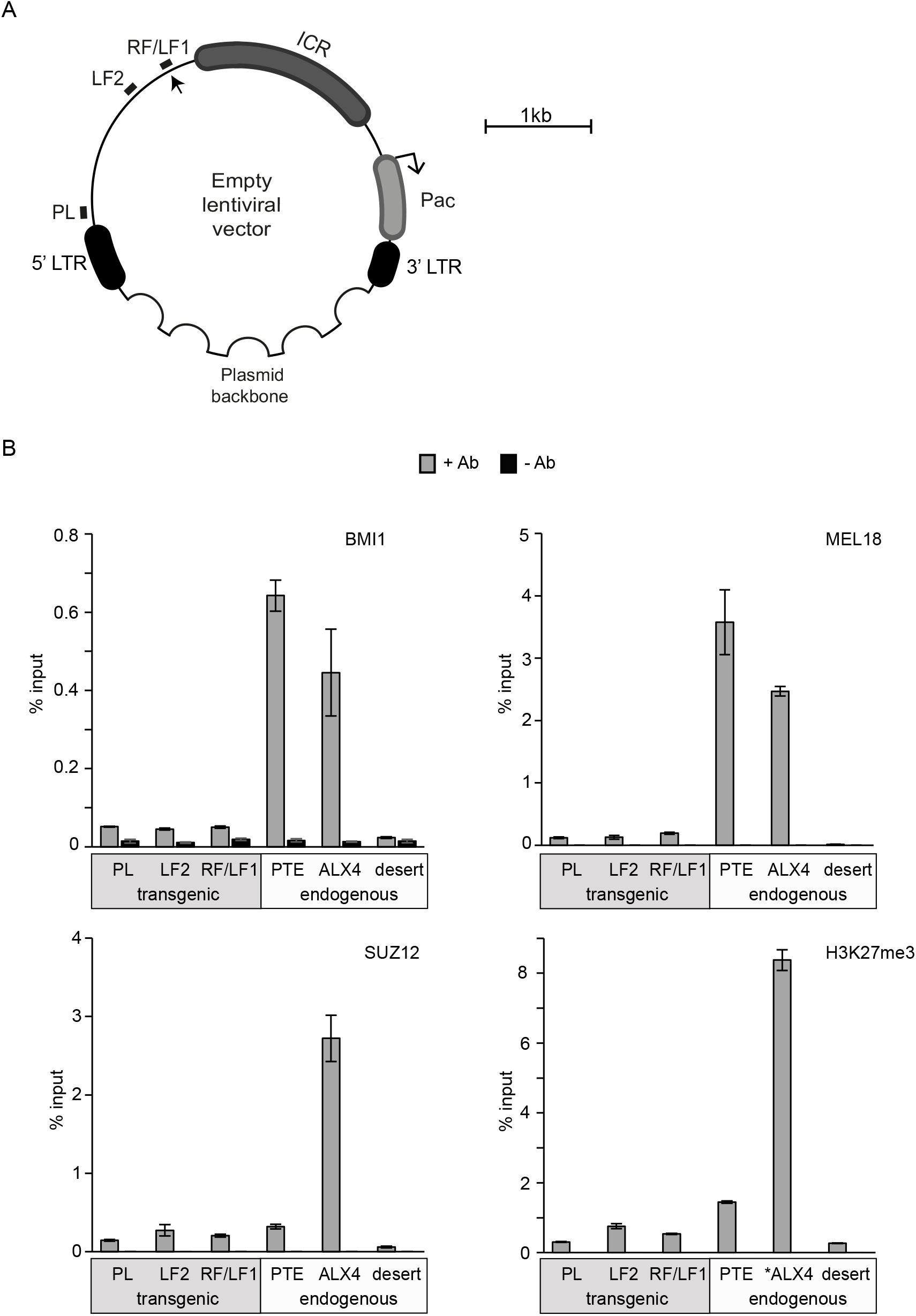
The empty lentiviral vector does not recruit PcG complexes. **A.** The schematic of lentiviral vector backbone. The vector contains LTR sequences required for genomic integration and *Pac* puromycin resistance gene flanked by the chromatin insulator from mouse *Igf2/H19* Imprinting Control Region (ICR). Positions of amplicons used for ChIP-qPCR analyses in **B.** are indicated as short lines. Note that LF1/RF amplicon overlaps the restriction site (arrowhead) used for cloning of PTE and control fragments. **B.** The ChIP-qPCR analyses with the indicated antibodies show the background binding that is much weaker than that seen at the endogenous *CCND2* PTE or *ALX4* gene. The histograms show the average and the scatter (whiskers) between the two independent experiments.

**Figure S7.**
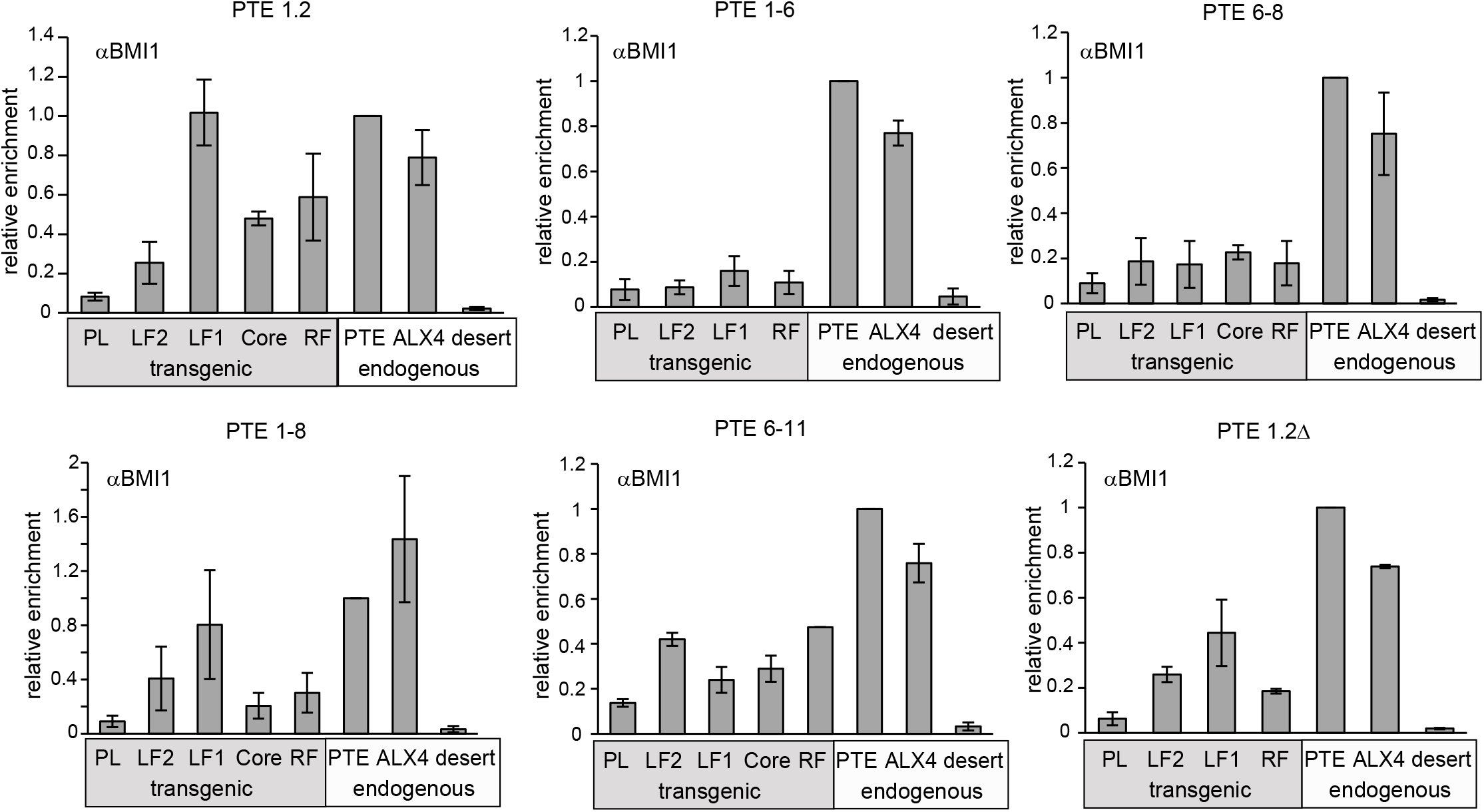
ChIP analysis of the BMI1 recruitment by PTE 1.2 sub-fragments. Positions of various amplicons along the transgenic constructs (grey box) are as in Figure 3A. Note that some transgenic constructs lack some of the amplicons. Precipitation of transgenic fragments is compared to internal positive and negative controls (white box). The histograms show the average and the scatter (whiskers) between the two independent experiments.

**Figure S8.**
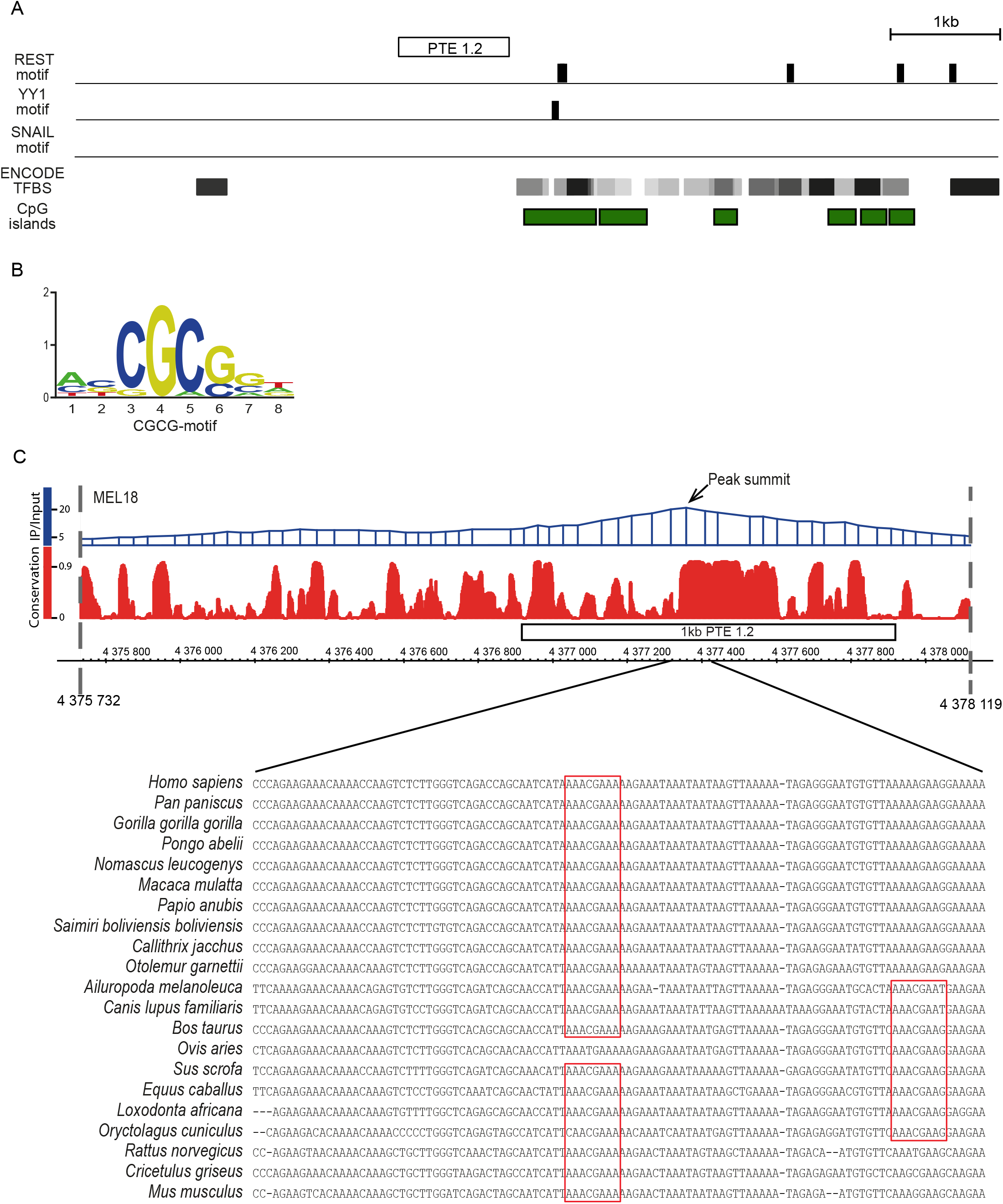
Distribution of sequence motifs in the vicinity of the *CCND2* PTE. **A.** The top tracks show the distribution of predicted recognition sequences for YY1, REST and SNAIL proteins. A composite distribution of all transcription factors (TFBS) mapped by ENCODE and positions of predicted CpG-islands (green rectangles) are shown below. **B.** Logo representation of the “CGCG”-motif. **C.** The alignment of DNA sequences from 21 vertebrate species and PhastCons score (lower panel) illustrate that the sequence of “CGA”-motif (red boxes) within *CCND2* PTE is highly evolutionary conserved. The majority of the species have one “CGA”-motif in the same position as in the human sequence. Some species have the second instance of the “CGA”-motif. In sheep (*Ovis aries*) the first instance of the motif is absent but the second one is present.

**Figure S9.**
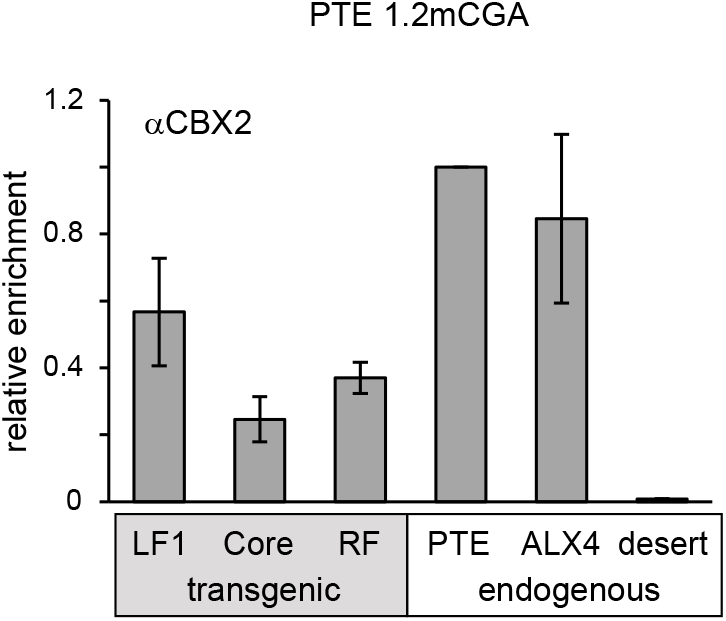
ChIP analysis of the CBX2 binding to the PTE 1.2mCGA transgene. Precipitation of transgenic fragment (grey box) is compared to internal positive and negative controls (white box). Positions of amplicons within the construct are indicated in Figure 3A. The histograms show the average and the scatter (whiskers) between the two independent experiments.

**Figure S10.**
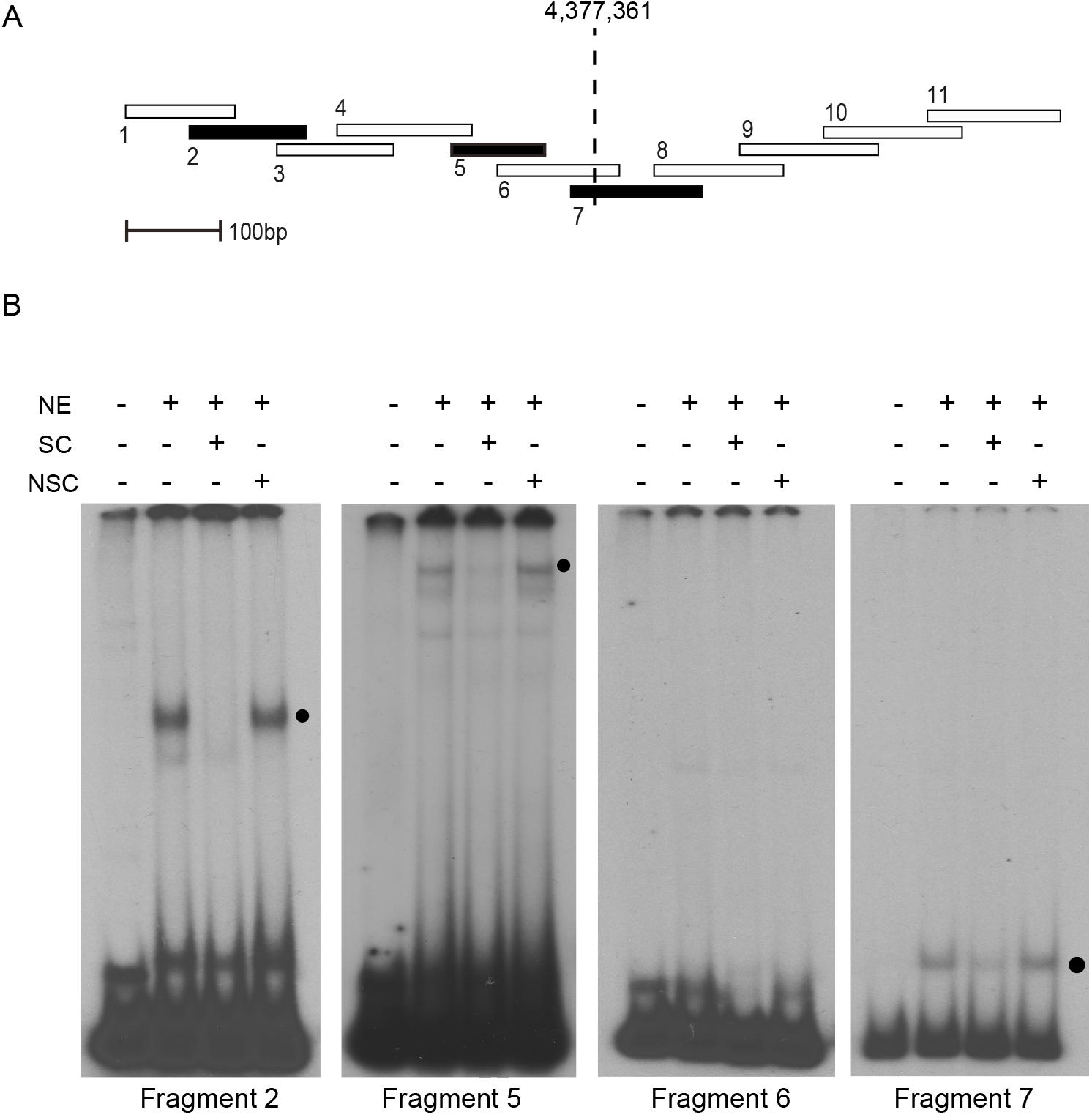
Sequence specific binding of nuclear proteins to the 1kb PTE 1.2. **A.** 100-150bp DNA fragments tiling 1kb PTE 1.2 are shown as rectangles. Fragments that show *in vitro* binding are marked in black. The vertical dashed line indicates the position of the *CCND2* MEL18/BMI1 binding peak summit. **B.** Representative EMSA experiments demonstrate that the fragments 2, 5 and 7 but not the fragment 6 shift the electrophoretic mobility (black circles) after incubation with the nuclear extract (NE). This shift is inhibited by the addition of an excess of specific unlabeled competitor (SC) while the addition of nonspecific competitor (NSC) has no effect. In this example fragment 5 was used as a nonspecific competitor for the EMSA with fragment 2 and vice versa. This illustrates that the fragments 2 and 5 are targeted by distinct sequence specific DNA binding activities. Similar tests were done for the fragment 7 (not shown). Note that the image of the EMSA with fragment 7 was acquired after much shorter exposure than those for fragments 2, 5 and 6.

**Figure S11.**
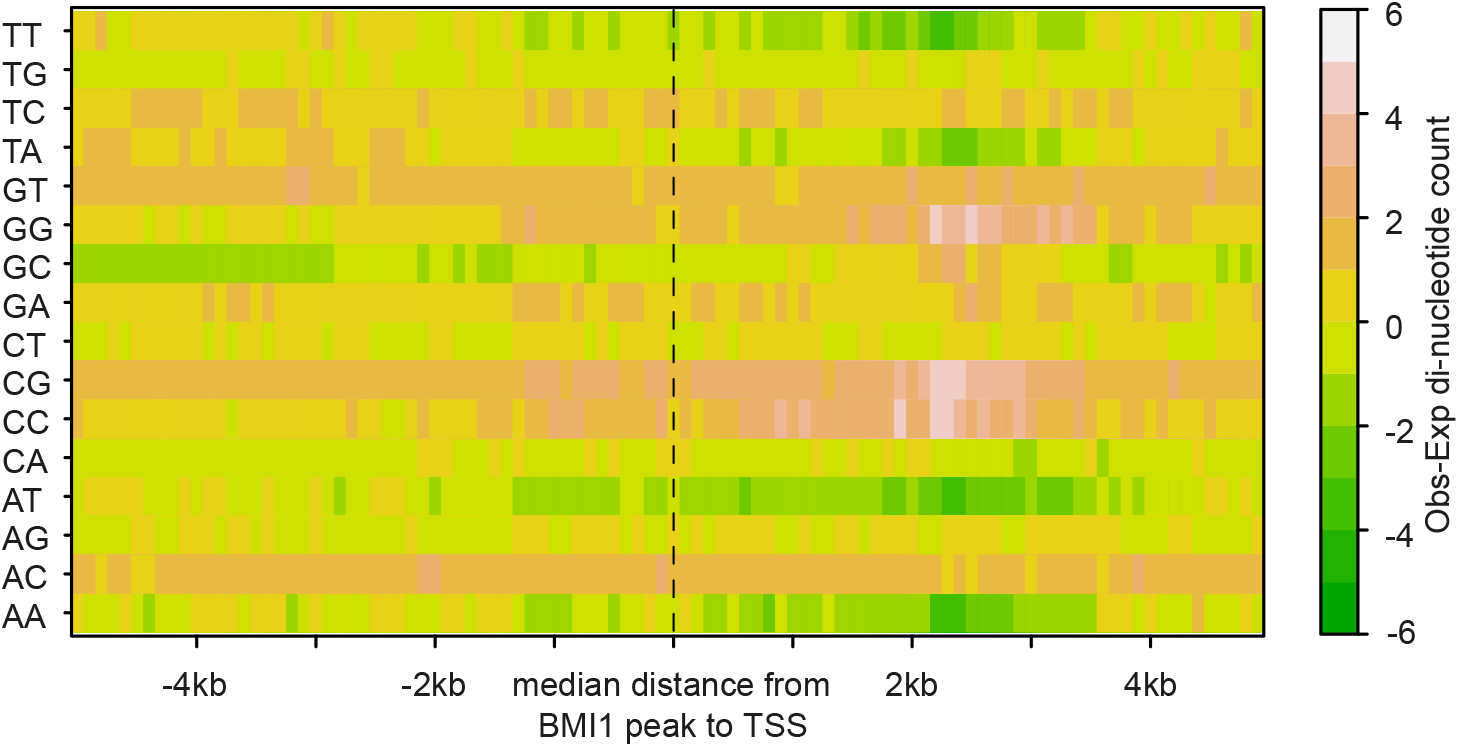
The control heat-map of di-nucleotide distributions within 5 prime regions of human genes. Each square represents the difference between the mean di-nucleotide count within a 100 bp window (observed) and the average count throughout the genome (expected). The midpoint (dashed line) corresponds to position +2355bp upstream of all annotated human TSS, which matches the median distance from a high-confidence BMI1 peak to the nearest TSS.

**Figure S12.**
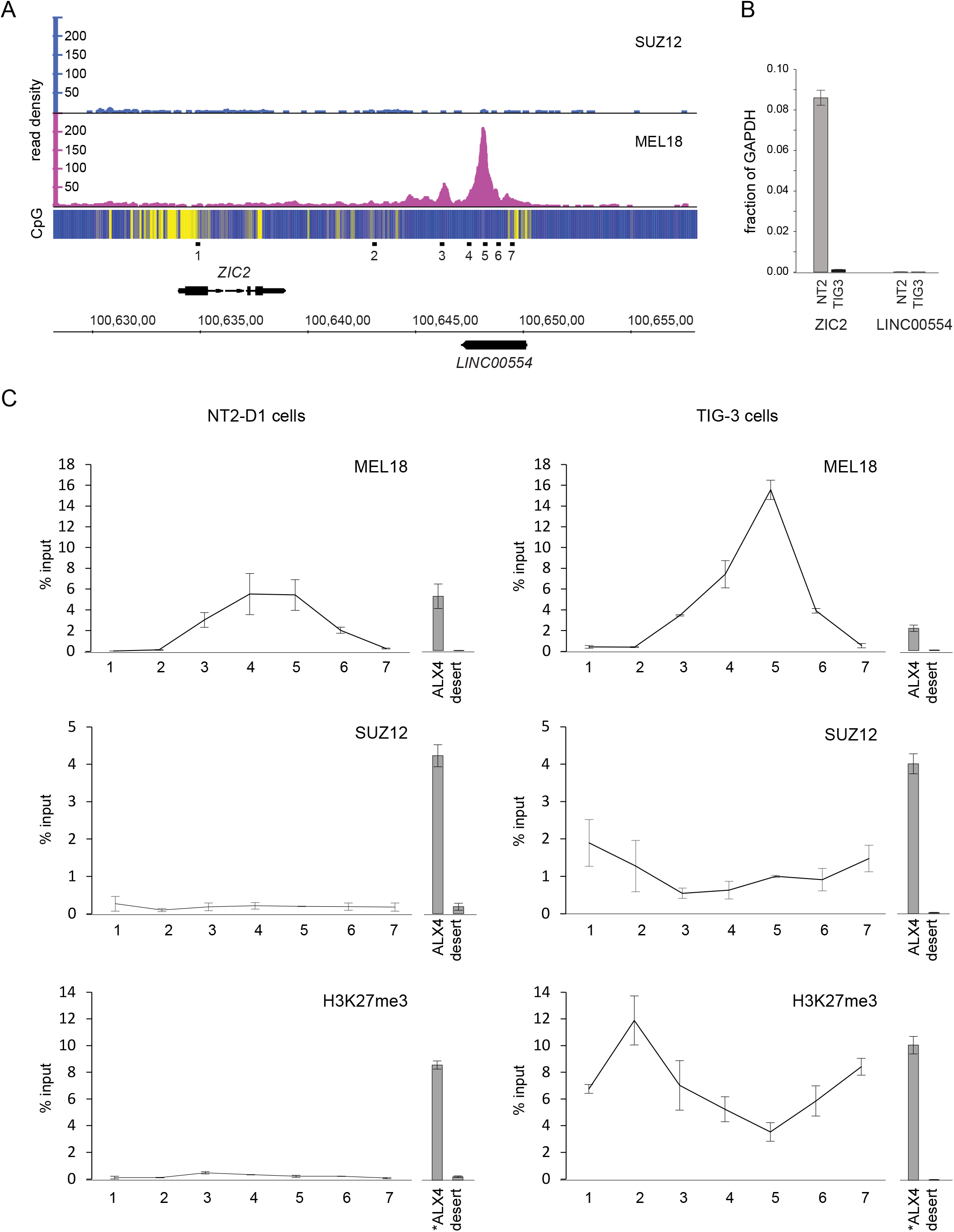
*ZIC2* gene contains a putative PTE. **A.** The schematic of human *ZIC2* locus. ChIP-seq with chromatin from NT2-D1 cells indicates strong binding of MEL18 but not SUZ12 to a narrow region (a putative PTE) downstream of *ZIC2* gene. The heat-map underneath ChIP-seq profiles shows the number of CpG nucleotides within 100bp sliding window (ranging from dark blue=0 to bright yellow=10). The *ZIC2* gene is transcribed from left to right and the lncRNA *LINC00554* of unknown function is transcribed from right to left. Positions of PCR amplicons analyzed in **B** and **C** are shown below the scale in Hg19 genomic coordinates. **B.** RT-qPCR measurements indicate that *ZIC2* is highly expressed in NT2-D1 (NT2) but very little in TIG-3 cells. *LINC00554* is not expressed in either of the cell lines. Histograms show the average and the scatter (whiskers) between two independent experiments. Values are normalized to the expression of the housekeeping *GAPDH* gene. **C.** ChIP profiles of MEL18, SUZ12 and H3K27me3 in NT2-D1 (left column) and TIG-3 (right column) cells. The immunoprecipitation of ALX4 gene repressed by PcG in NT2-D1 and TIG-3 cells was used as positive control and Chromosome 12 gene desert was assayed as negative control. Histograms for both are shown to the right of the each graph and are to the same y-axes scale. ALX4* positive control region for H3K27me3 ChIPs is offset from the ALX4 region used for the profiling of other proteins. All histograms and graphs show the average and the scatter (whiskers) between two independent experiments. Note that in TIG-3 cells H3K27me3 and Suz12 profiles are offset from the MEL18 peak into neighboring CpG-rich regions.

**Table S1.**
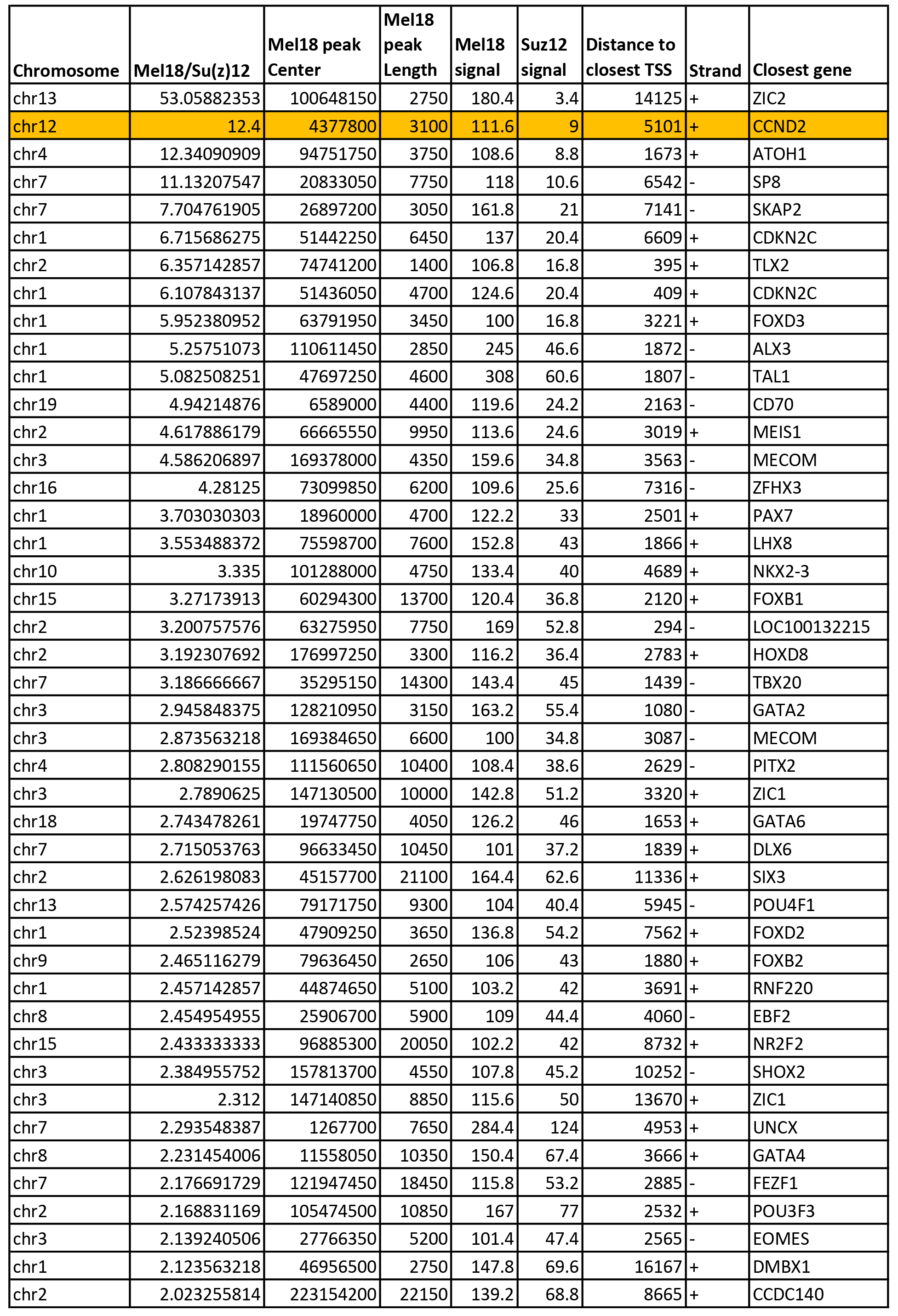
Screening for MEL18 sites with weak binding of SUZ12. Coordinates are in genomic release Hg19.

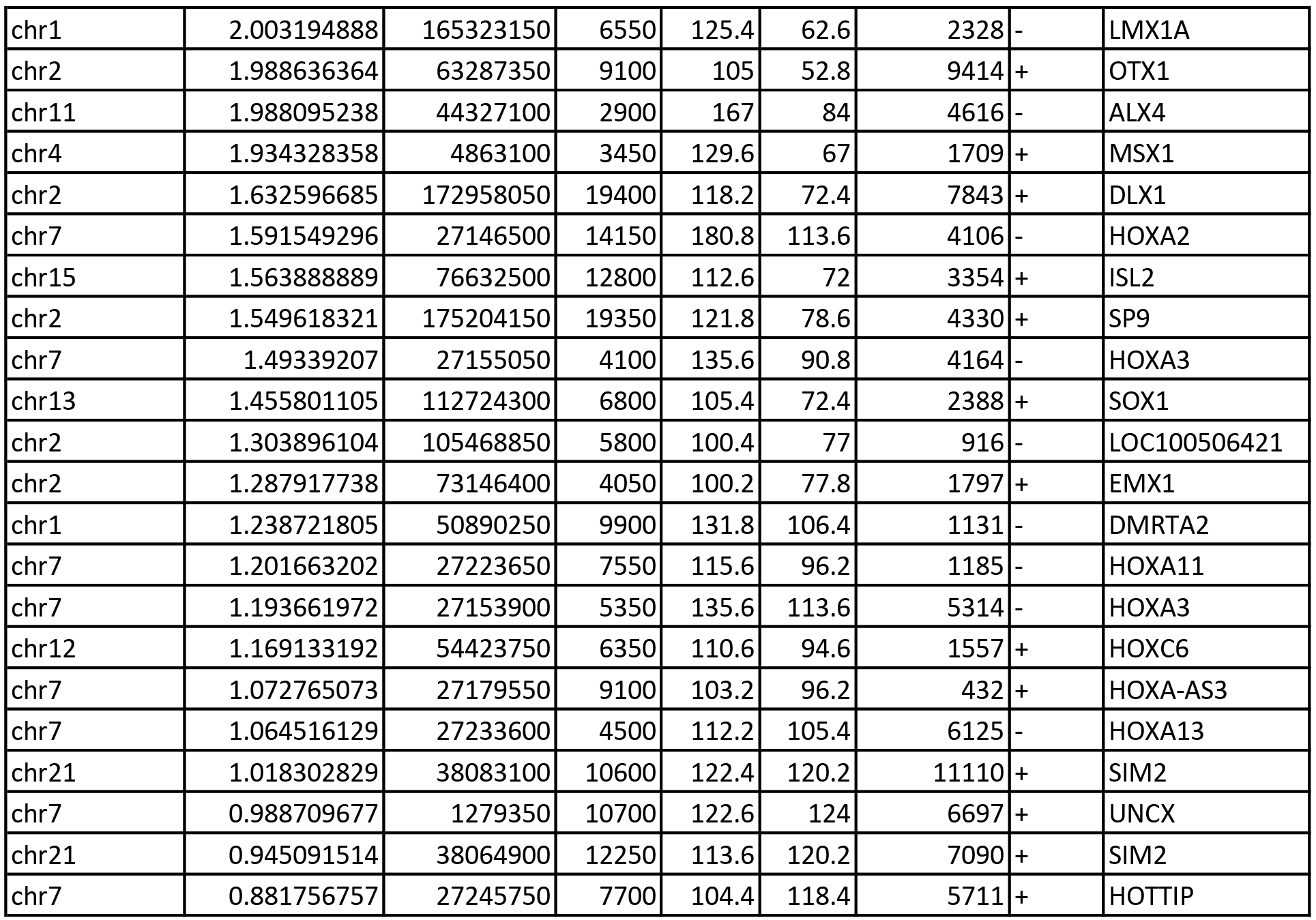

**Table S2.**
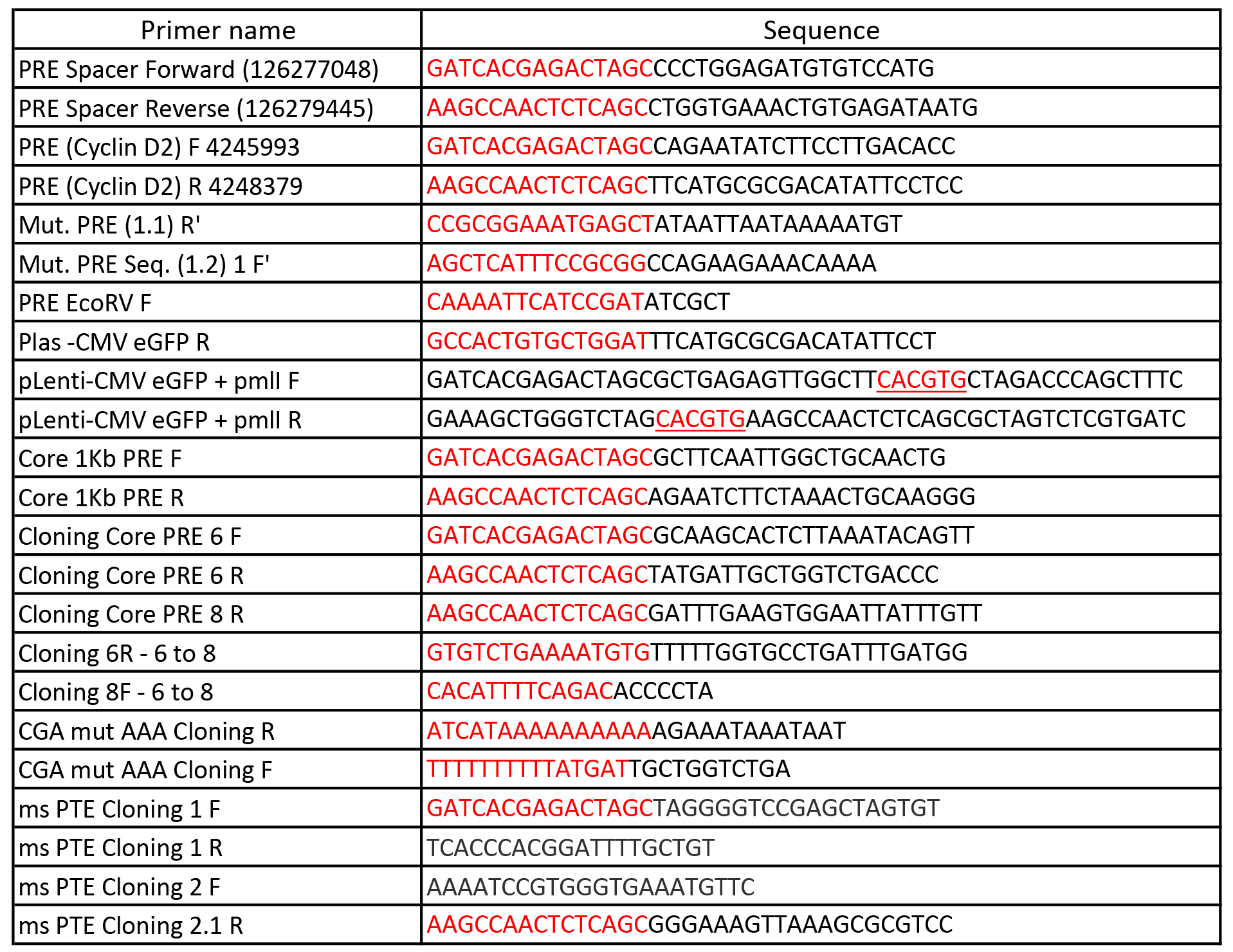
The sequence and name of PCR primers for cloning.

The nucleotides in red are the 15nt homology sequence required for In-Fusion cloning. Underlined nucleotides indicate introduced restriction sites.

**Table S3.** The correspondence between PCR primers and transgenic constructs.

| construct | Primer name |
| --- | --- |
| gene desert | PRE Spacer Forward (126277048) |
|  | PRE Spacer Reverse (126279445) |
| PTE 2.4 | PRE (Cyclin D2) F 4245993 |
|  | Mut. PRE (1.1) R' |
|  | Mut. PRE Seq. (1.2) 1 F' |
|  | PRE (Cyclin D2) R 4248379 |
| PTE 1.1 | PRE (Cyclin D2) F 4245993 |
|  | Mut. PRE (1.1) R' |
| PTE 1.2 | Core 1Kb PRE F |
|  | Core 1Kb PRE R |
| PTE 1.3 | Core 1Kb PRE F |
|  | PRE (Cyclin D2) R 4248379 |
| PTE 1-6 | Core 1Kb PRE F |
|  | Cloning Core PRE 6 R |
| PTE 1-8 | Core 1Kb PRE F |
|  | Cloning Core PRE 8 R |
| PTE 6-8 | Cloning Core PRE 6 F |
|  | Cloning Core PRE 8 R |
| PTE 6-11 | Cloning Core PRE 6 F |
|  | Core 1Kb PRE R |
| PTE 1.2Δ | Core 1Kb PRE F |
|  | Cloning 6R - 6 to 8 |
|  | Cloning 8F - 6 to 8 |
|  | Core 1Kb PRE R |
| PTE 1.2 CGA mut | Core 1Kb PRE F |
|  | CGA mut AAA Cloning R |
|  | CGA mut AAA Cloning F |
|  | Core 1Kb PRE R |
| Mouse PTE | ms PTE Cloning 1 F |
|  | ms PTE Cloning 1 R |
|  | ms PTE Cloning 2 F |
|  | ms PTE Cloning 2.1 R |

**Table S4.**
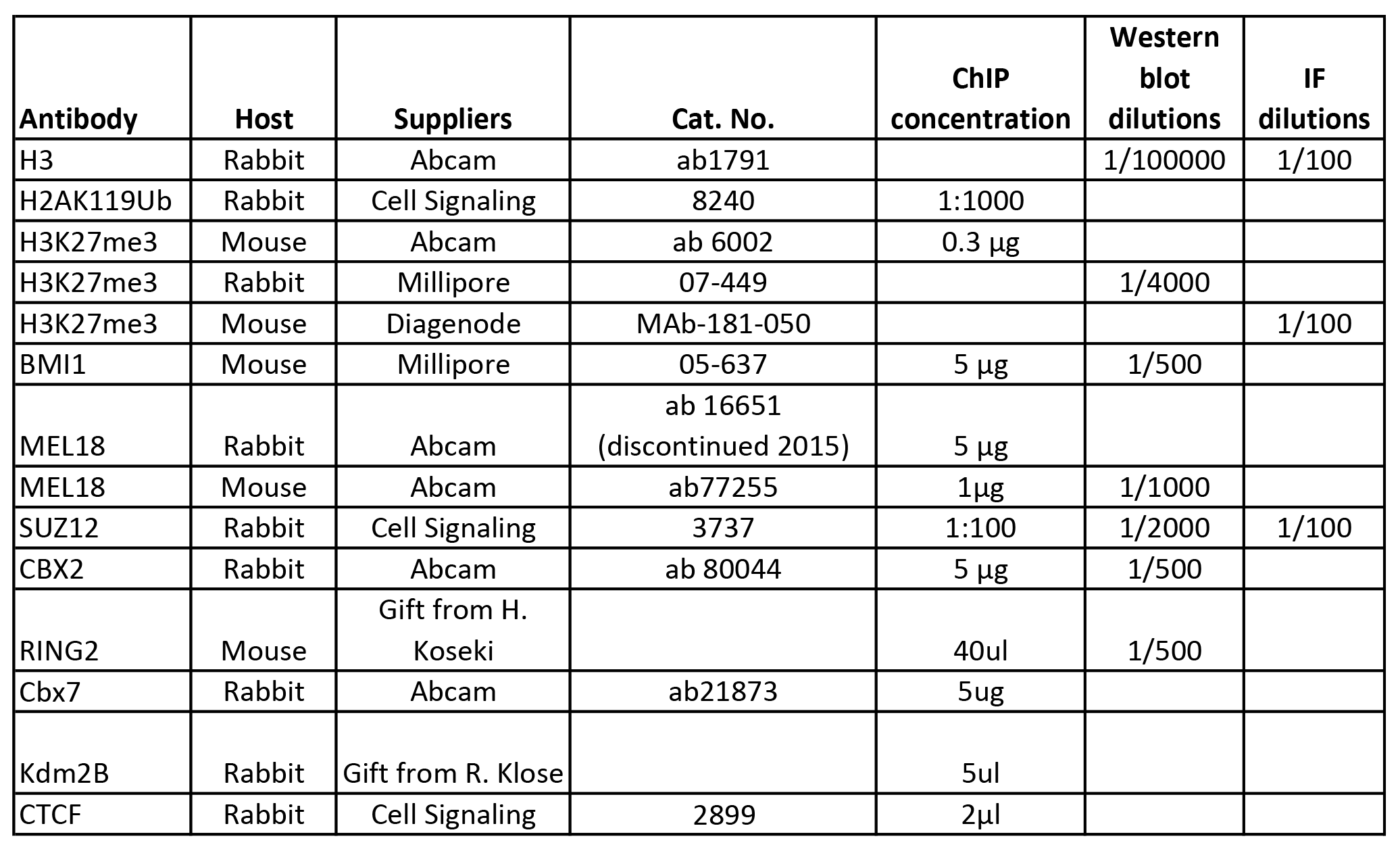
List of antibodies used for ChIP analysis, western blot and immunofluorescence.

**Table S5.**
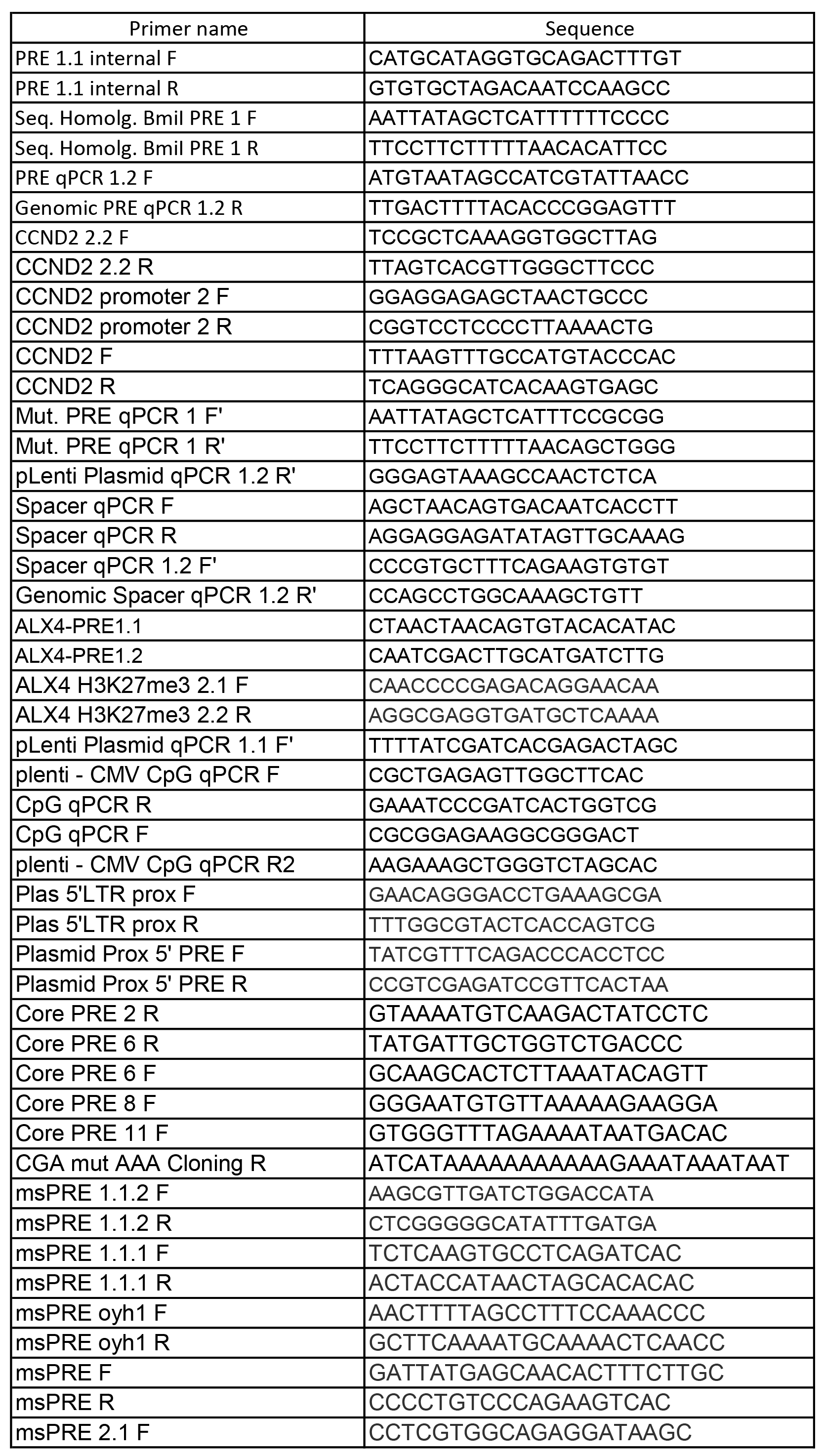
The PCR primers used for ChIP-qPCR, RT-qPCR and genotyping.

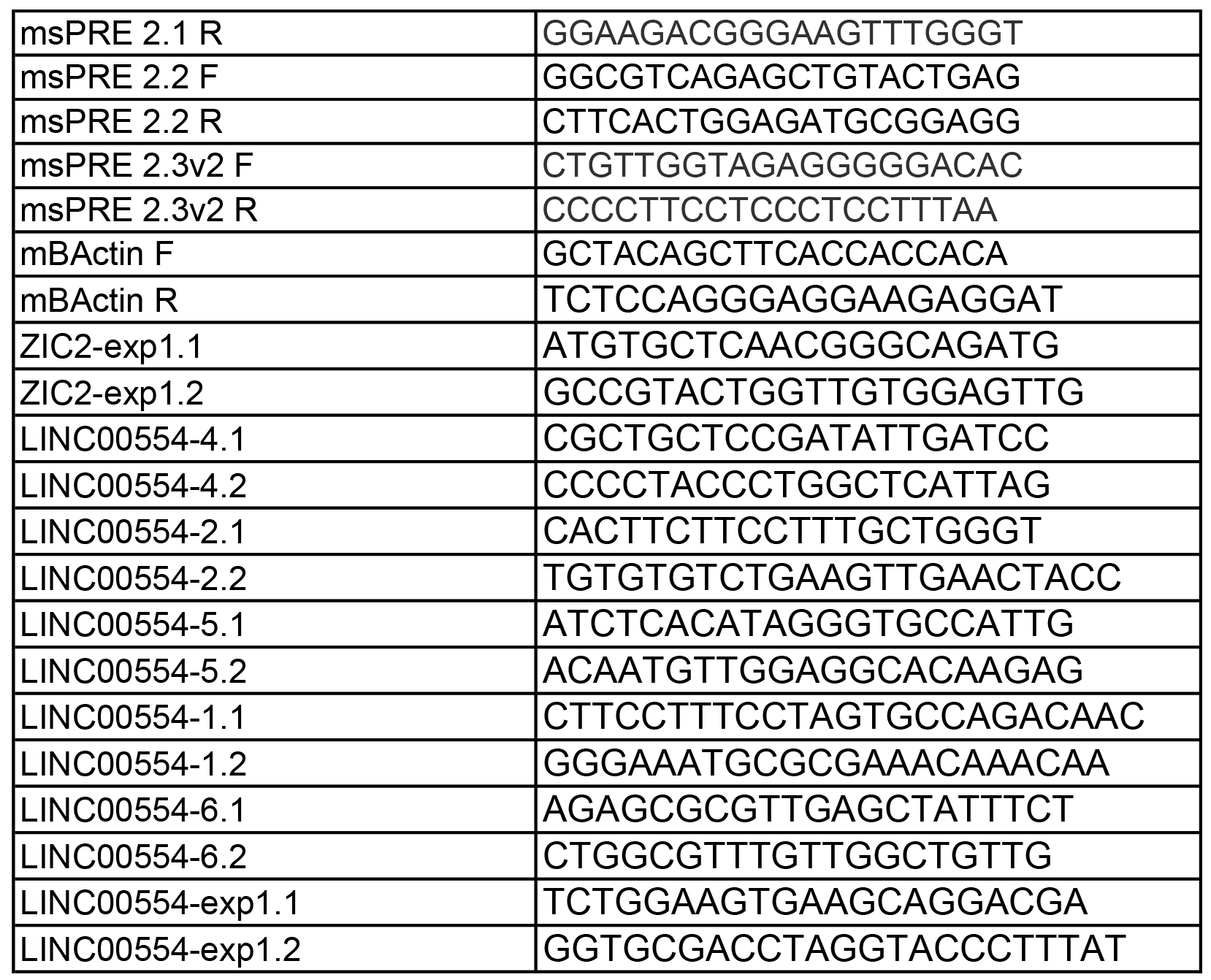

**Table S6.**
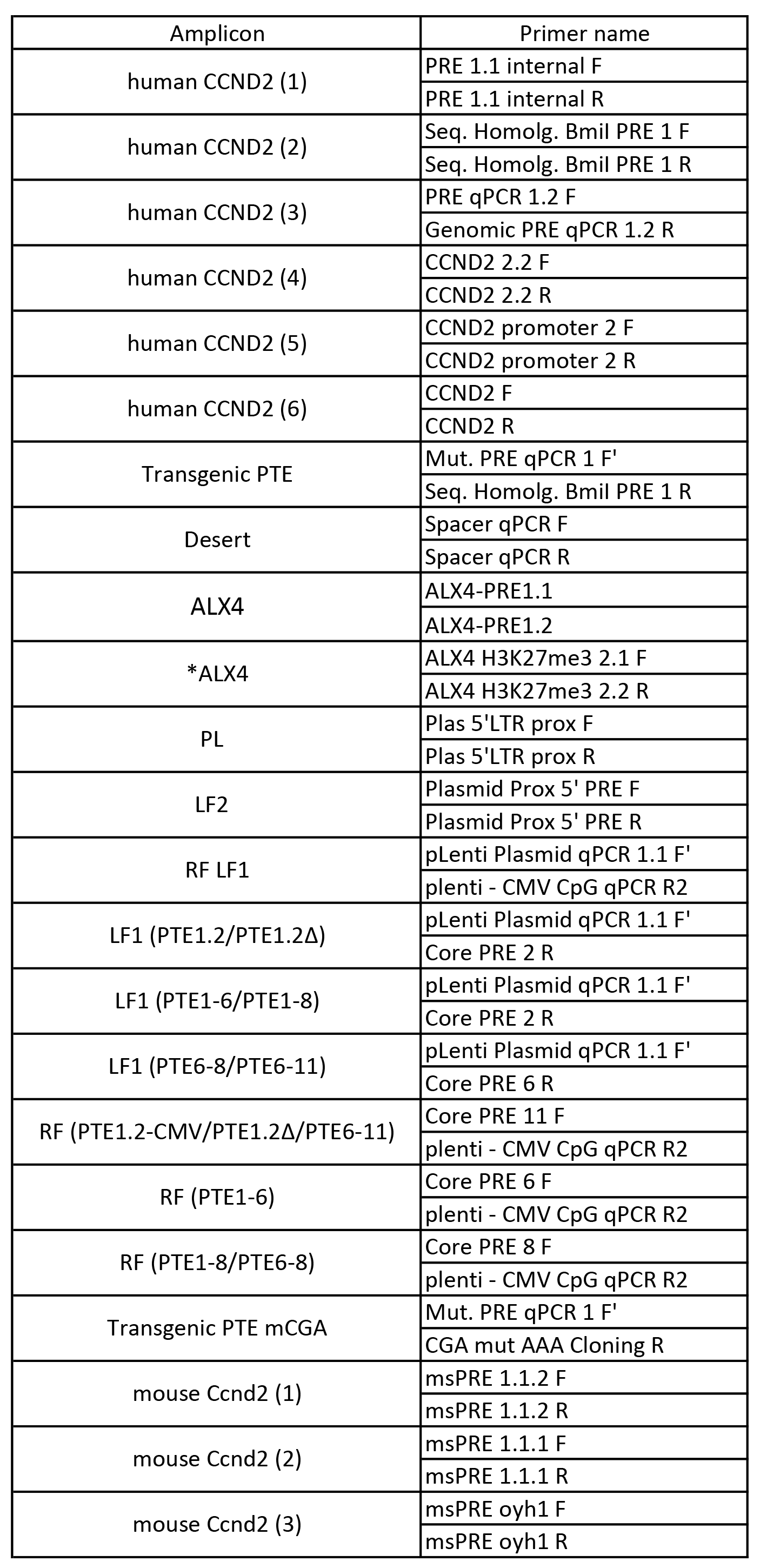
The correspondence between combination of PCR primers and amplicons.

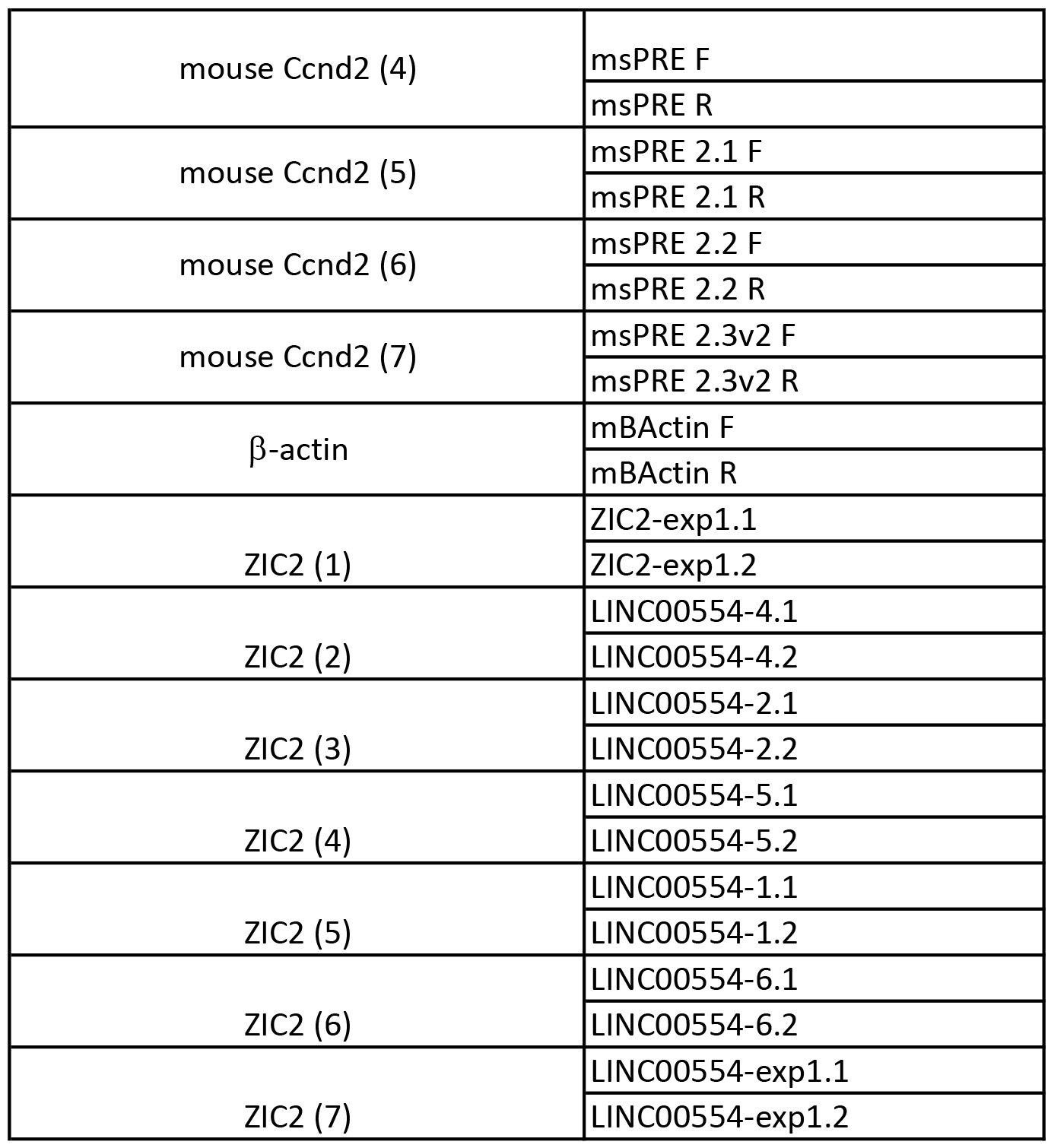

**Table S7.**
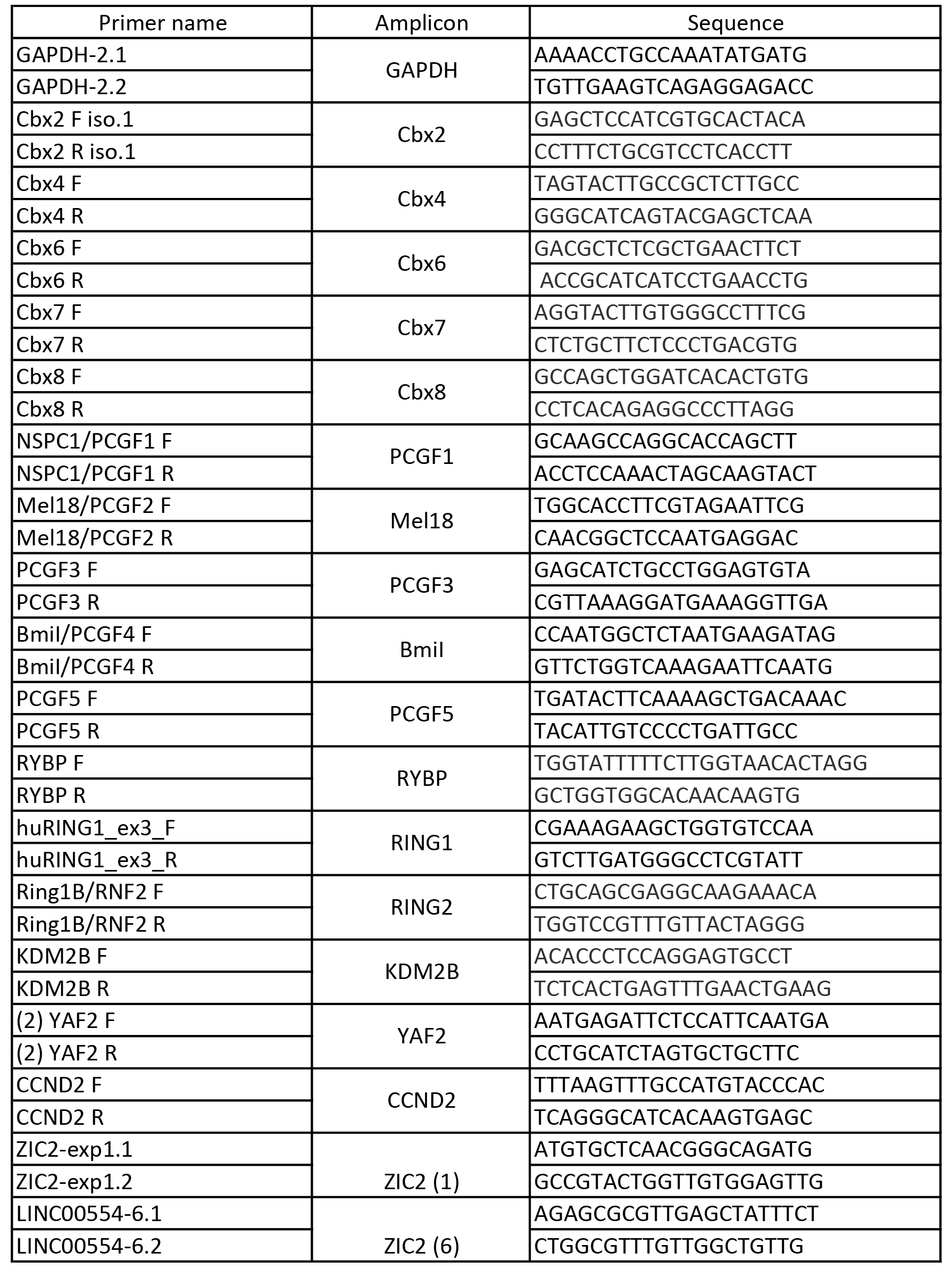
The PCR primers used for RT-qPCR analysis.

**Table S8.**
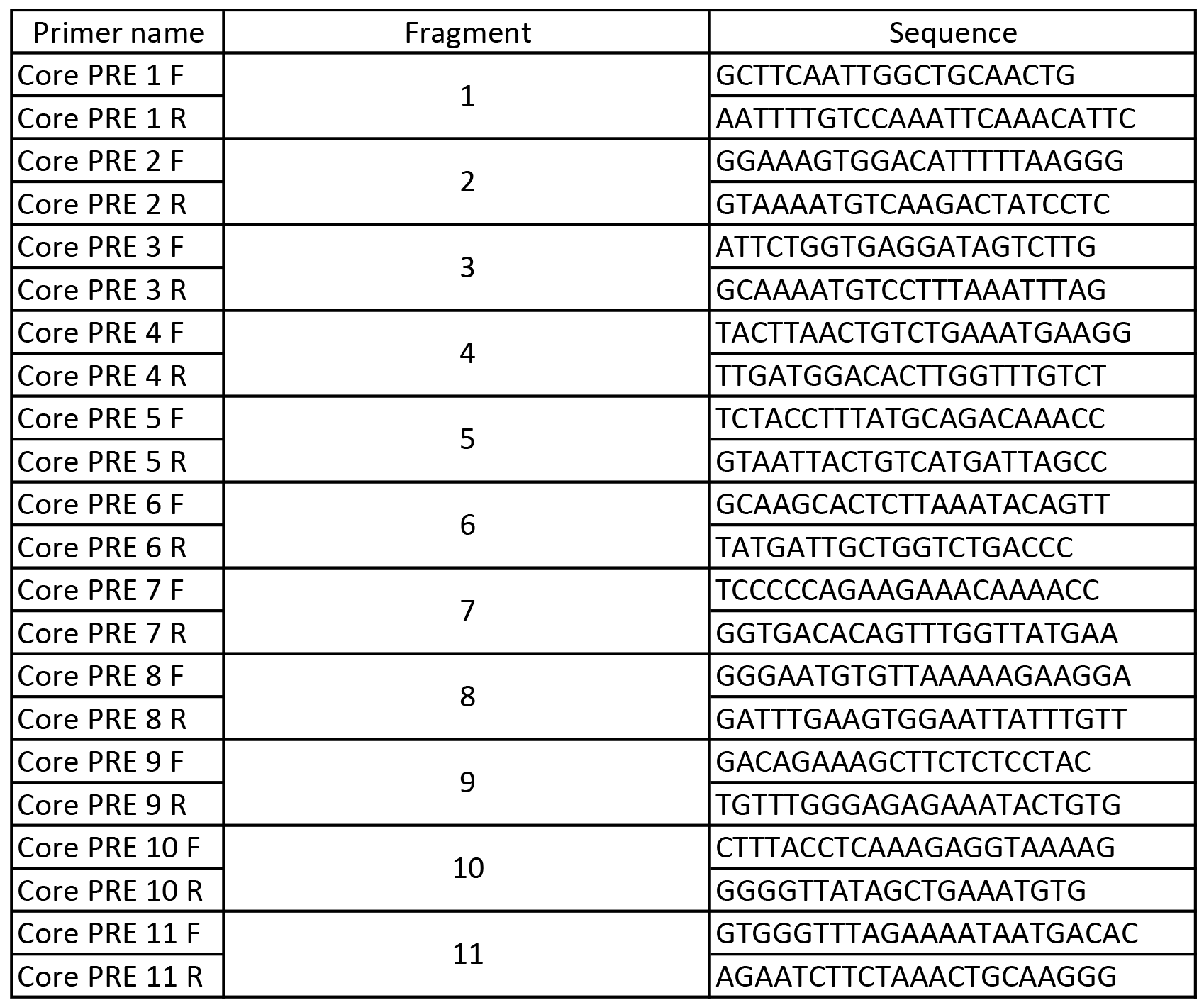
PCR primers to produce and label the fragments for EMSA.

